# TIMS-Bench: Towards community standards for benchmarking untargeted trapped ion mobility metabolomics tools and datasets

**DOI:** 10.64898/2026.05.23.724673

**Authors:** Prajit Rajkumar, Yojana Gadiya, Victoria Deleray, Aurelie Roux, Kiana A West, August Allen, Pieter C Dorrestein, Daniel Domingo-Fernández, Biswapriya B Misra

## Abstract

Untargeted liquid chromatography–tandem mass spectrometry (LC-MS/MS)-based metabolomics is an important technology for unbiased discovery of small molecules in biomedical (e.g., drug discovery to diagnostics), animal, plant, environmental, and microbial research. Over the past decade, ion mobility has added an additional dimension to the triplet of MS1, MS2, and retention time, helping resolve co-eluting or isomeric features in an LC-MS/MS that aid in compound identification. Here, we focused on evaluating the current trapped ion mobility spectrometry (TIMS)-amenable feature-finding tools (MZmine 4.9, MS-DIAL 5.5, and MetaboScape 2025 14.0.3) for pre-processing of metabolomics data generated using a popular ion mobility mass spectrometry (IM-MS) technique, TIMS. We leveraged ten public and three benchmark TIMS datasets to evaluate these tools for their strengths and weaknesses. Our results show that MZmine consistently identified the highest number of features and confidently annotated features; however, this performance was accompanied by an increased number of false positives, due to peak splitting, as well as reduced accuracy in collision cross section (CCS) measurements. In contrast, MetaboScape achieved the highest fraction of high-quality MS2 spectra, reflecting a more conservative feature detection strategy. MS-DIAL demonstrated balanced performance, identifying features that other tools missed. Finally, we publicly release the ground-truth datasets and code to support future developments in improving IMS data analysis.

## INTRODUCTION

The advent and application of ion mobility mass spectrometry (IM-MS) in the omics era have led to adoption across proteomics, metabolomics, lipidomics, exposomics, and glycomics research^1^. In addition to the limited MS1, MS2, and retention time (RT) that enable the unambiguous identification of compounds with standard compounds and/or reference libraries, IM-MS-driven ion mobility (i.e., collision cross-section (CCS) values) offers an orthogonal fourth dimension. This additional dimension enhances confidence in metabolite annotation rates while enabling the resolution of complex isomers beyond chromatography and mass-to-charge (*m/z*) dimensions, but challenges remain in terms of extracting the data^2^.

Metabolomics researchers have long recognized that mass accuracy alone was insufficient for metabolite identification, necessitating the addition of orthogonal dimensions, such as chromatographic separation and IM-MS. IM-MS utilizes rapid gas-phase separation to enhance peak capacity and signal-to-noise (S/N) ratio, adding the ion mobility dimension (i.e., CCS values) as an additional molecular descriptor. This approach mitigates issues like matrix interference and long run times^3^, providing an additional data layer for metabolite annotation by ruling out other candidate molecules that do not share the same drift times^4^. IM-MS also enhances metabolome coverage and throughput by resolving isobaric/isomeric metabolites that co-elute in LC^2^, thereby improving resolution and sensitivity. Consequently, IM-MS is applied in hybrid workflows such as direct infusion IM-MS and MS imaging^5^. Users now have access to several variants of IM-MS instruments from various vendors, including drift tube ion mobility spectrometry (DTIMS), traveling-wave ion mobility spectrometry (TWIMS), trapped ion mobility spectrometry (TIMS), field□asymmetric waveform ion mobility spectrometry (FAIMS), and cyclic IMS. TIMS can provide accurate CCS values (<0.2% RSD) and a high mobility resolving power (*R* up to 470)^6^.

Alongside data generation, data analysis has emerged as a specialized and critical component of omics research. Dozens of processing tools are available for analyzing untargeted high-resolution mass spectrometry datasets, such as MZmine 3^7^, MS-DIAL^8^, Met4DX^9^, and xcm^10^. Only the former three have been reported to handle IM-MS datasets. In addition, proprietary tools, such as MetaboScape (Bruker Daltonics), available to a TIMS-TOF user, enable competent processing of TIMS data while retaining the IM dimension intact. To the best of our knowledge, at least 20 publications report the evaluation of existing tools on non-TIMS untargeted metabolomics data. However, the majority of these publications focus on the development of novel software **(Supplementary File 1)**, in part because the high cost of time and resources required for manual annotation has hindered the establishment of benchmarking or ground truth datasets in metabolomics. Consequently, researchers must rely on *de novo* generation of ground truth data, in which compound identities and abundances are predefined in a background matrix before untargeted LC-MS/MS acquisition. Despite the increasing adoption of TIMS datasets, only one prior study has evaluated untargeted metabolomics tools for this technology. That work, however, was restricted to human and mouse samples and reported only feature counts and overlaps. Such metrics are insufficient for assessing performance, as a higher feature count may reflect noise or artifacts rather than true positives. Furthermore, because the authors evaluated their own software, Met4DX, the study may be geared towards testing the performance of Met4DX specifically, as evidenced by the restricted feature overlap analysis, which computes pairwise feature overlaps for Met4DX against MZmine 3, MS-DIAL, and Metaboscape, but does not compute overlaps between pairs of tools that do not include Met4DX^9^.

In this work, we present one of the first evaluations of untargeted metabolomics tools on IM-MS data using benchmark data. We assess two open-source available tools, MZmine 4.9^7^and MS-DIAL 5.5^11^, as well as the proprietary software, MetaboScape 2025 14.0.3 (Bruker Daltonics). The evaluation is performed using three ground-truth datasets, two of which we generated, and one that is publicly available, as well as ten publicly available datasets that we identified. Collectively, these 13 datasets encompass a range of species and modalities, enabling a comprehensive evaluation of the advantages and limitations of the tools described above. We evaluate tools across a broad range of tasks, spanning from basic feature-level metrics (e.g., the total number of features, the number of MS2-annotated features, and the accuracy of CCS values) to performance-related metrics (e.g., precision and recall) on ground-truth datasets. Overall, our findings inform and guide the community by highlighting the strengths of existing tools while fostering ideas for developing new ones.

## MATERIALS AND METHODS

### Datasets

In this evaluation, we employed 10 publicly available timsTOF datasets **(Figure 1)**, acquired using TIMS technology, specifically 4D metabolomics with ion mobility (IM) mode ON on the timsTOF instruments from Bruker Daltonics **(Supplementary Table 1)**^7,12-16^. We carefully selected public datasets covering a wide range of biological matrices, including plant extracts, animal tissues, environmental samples, microbial samples, and cell lines. Furthermore, we evaluated two internally generated and now publicly available ground-truth datasets as described below.

**Figure 1.**
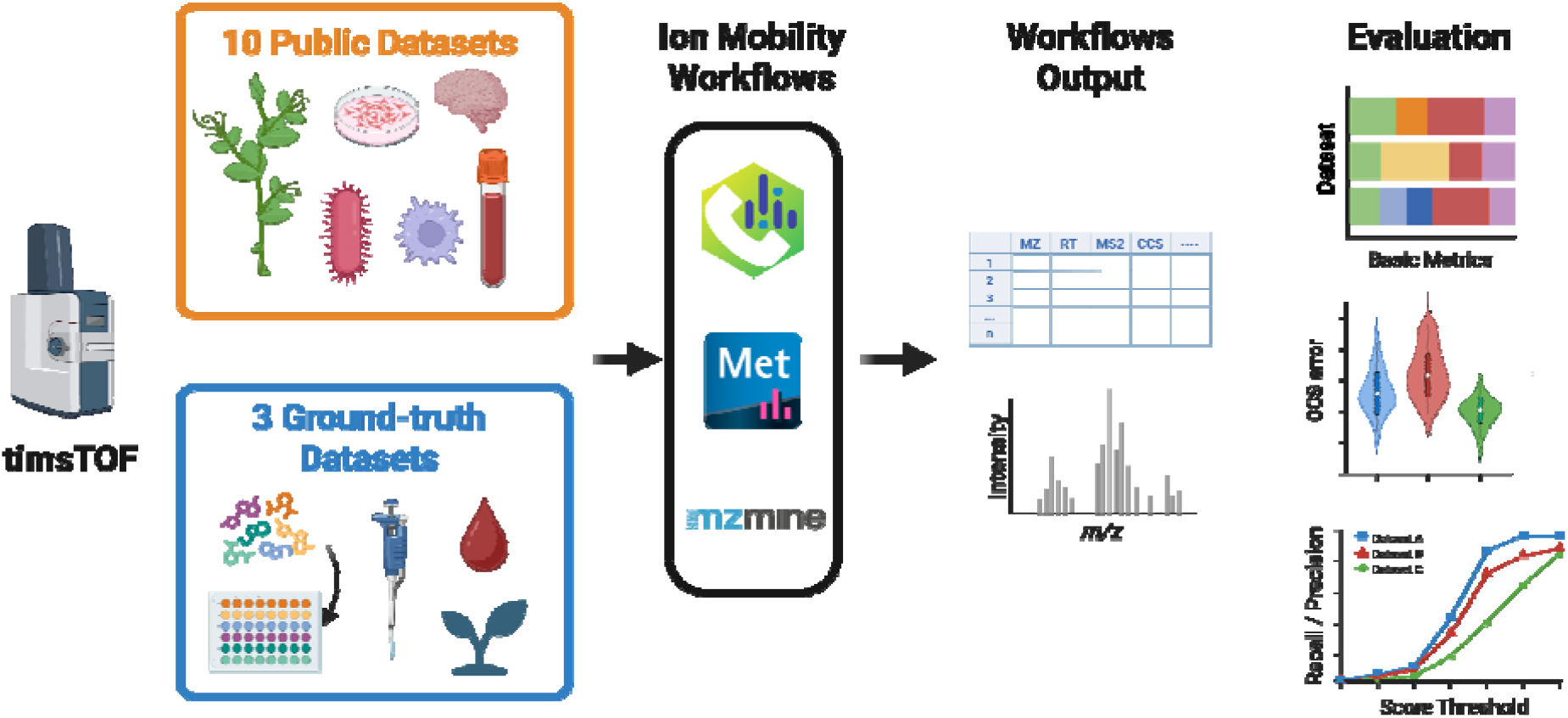
Schematic illustration of the evaluation. Created with BioRender.com.

### Ground-truth datasets

#### ReFRAME library dataset

The first ground-truth dataset comes from an ongoing study in the Skaggs School of Pharmacy and Pharmaceutical Sciences at University of California, San Diego, conducted by Pieter Dorrestein’s group, which comprises pools of 4,723 drug standards that are part of an expansion to empirically perform medication readouts from untargeted metabolomics data^17^. These mixtures underwent LC-TIMS-PASEF analysis in a Bruker timsTOF Pro2 coupled to an Agilent 1260 Infinity III LC to produce a spectral and CCS library (MassIVE accession: MSV000098263). These drugs were a subset of a larger ReFRAME (Repurposing, Focused Rescue, and Accelerated Medchem) library^18^. We refer to this dataset as “ReFRAME library dataset” throughout the manuscript.

#### Spiked-in NIST SRM 1950 human blood plasma dataset

We also generated another ground-truth dataset specifically for this investigation to provide a more challenging evaluation, in which we spiked NIST SRM 1950 human plasma (National Institute of Standards and Technology, USA), a widely used reference for analyte detection within a biological matrix, with a mixture of 126 additional metabolites spanning six biochemical classes (**Supplementary File 1)**. To measure the sensitivity of the tools for feature picking, linearity, and dynamic range, we used four concentrations (low, medium, high, and super-high) of the spiked standards. The resulting plasma aliquots were extracted by methanol precipitation, dried, and reconstituted prior to analysis as described (**Supplementary Text 1**). We collected data from 5 sample groups (mixtures of 126 compounds spiked at (i) low, (ii) medium, (iii) high, (iv) super-high doses in plasma and (v) non-spiked plasma) with technical replicates (n = 8). We refer to this dataset as the “spiked plasma dataset” throughout the manuscript.

#### Plant natural products dataset

To test the three tools’ ability to detect a diverse set of compounds, we generated a second ground-truth dataset for this study. This dataset comprised 45 plant-derived compounds (MW ranging from 124.14 - 504.7 Da and belonging to natural product classes such as alkaloids, phenylpropanoids, phenolics, flavonoids, triterpenoids, among others) **(Supplementary Table 2)**, which were procured commercially and run individually at three different on column amounts (50, 100, and 200 ng) (**Supplementary Text 2)**. We refer to this dataset as the “plant natural products dataset” throughout the manuscript.

For both the NIST SRM 1950 plasma and the 45-plant natural products dataset, untargeted LC-MS/MS was performed in-house on a timsTOF Pro2 system using reversed-phase chromatography (C18 column) in positive ionization mode with 4D PASEF data acquisition. Detailed LC and MS conditions are provided (**Supplementary Text 3**).

#### Public datasets

To evaluate the tools against more challenging and diverse datasets, we leveraged publicly archived ten different timsTOF IMS (4D) datasets found across three repositories, Massive-GNPS^19^ (eight datasets), Metabolights^20^ (one dataset), and Metabolomics Workbench^21^ (one dataset). These public datasets span diverse biological origins, including plants, animals, microorganisms (two datasets each), standards and drugs, and humans (four datasets) **(Figure 1, Table S1)**. These public datasets were acquired across timsTOF flex (5 datasets) and timsTOF Pro/ Pro 2 (5 datasets). Four of these ten datasets were acquired in both positive and negative ionization modes, prompting us to limit the analysis to positive-ion mode data alone. Except for two datasets acquired on HILIC (i.e., MSV000091642 and MSV000096189), the remaining eight were acquired on C18 reverse-phase UPLC/UHPLC systems. These datasets also included two lipidomics studies, whereas the remaining eight were metabolomics, natural products, or drug screening datasets.

While collecting public datasets, we ensured that only 4D datasets available in the vendor’s (Bruker’s) format, .d files, were included, i.e., raw files that have not undergone any conversion. We excluded 3D datasets obtained on the vendor’s TOF instruments available in the repositories, as they did not acquire the ion mobility dimension of the data (i.e., they ran with TIMS mode OFF).

### Untargeted LC-IM-MS/MS metabolomics tools

In our benchmark, we leverage three main untargeted metabolomics tools that are compatible with ion mobility data **(Figure 1)**: MetaboScape 2025 14.0.3 (Bruker Daltonics, Bremen, Germany), MS-DIAL 5.5^11^, and MZmine 4.9^7^. For the vendor’s proprietary data analysis suite, MetaboScape, we used the feature-extraction and data-processing parameters presented in **Supplementary Text 4**. For MS-DIAL, we used the latest version (5.5.250221), which handles ion mobility data and allows for the direct processing of .d raw datasets on the graphical user interface (GUI) using the parameters described in **Supplementary Texts 5 and 6**. Lastly, for MZmine, we processed the raw data using the command line interface (CLI) version of Mzmine 4.9 with the parameters described in **Supplementary Text 7**.

In addition to the tool-specific settings, there are general settings that apply to all tools. For example, we used the same common adduct list across all three tools **(Supplementary Table 3)**. Calibration metadata were unavailable for these public datasets and were therefore not incorporated into the analysis. Given the diverse origins of public datasets, i.e., many studies not providing blank runs, and the minimal contribution of true features from blanks, we could not leverage blank subtraction as an option. In addition, given the study’s premise regarding confidence in feature calling and linking, and with very limited or no metadata available for many of these studies, we did not group the samples by biological origin or by any methodology or motivations described in the manuscripts associated with the data. Lastly, we obtained the feature tables from each tool and its MS2 peak lists, which were exported as .mgf and/or .csv files and further standardized for downstream processing **(Figure 1)**.

### Feature annotation

#### Spectral reference libraries

We annotated features obtained from all three tools using the following spectral libraries: NIST23’s HRAM tandem MS libraries (2.4M spectra of 51k compounds), MassBank of North America (MoNA) (MoNA, 2023), MassBank^22^, MassBank of Europe^23^, ReSpect (RIKEN tandem mass spectral database)^24^, MS^n^ library^25^, Critical Assessment of Small Molecule Identification (CASMI)^26^, and GNPS libraries^19^. We avoided using the built-in annotation modules within each tool, as our goal was to test the performance of feature calling and linking independently of tool-specific settings. To ensure consistent comparisons across the three tools, we standardized and applied the same downstream annotation process to all three. Lastly, all the metabolite annotations reported in this study correspond to Metabolomics Standards Initiative (MSI) Level 2 confidence, relying on matches to public and in-house MS/MS spectral libraries rather than validation with authentic standards^27^. Compounds were treated as annotated when associated with a unique InChIKey-14.

#### Spectral matching approaches

Prior to spectral matching, we applied a filter to all MS2s and removed fragments with a lower relative intensity with respect to the highest fragment than 0.01 (1%), as well as an absolute intensity equal to or lower than 10. This pre-filtering step, although common in the literature, is essential for a fair evaluation^28^. This is due to significant variation in the number of MS2 fragments produced across the different tools (as detailed in **Section 3.1** of the Results). We employed three distinct spectral matching approaches:

1. **Spectral entropy similarity**^29^. By default, a confident match has a similarity of 0.7 or higher, corresponding to a spectral entropy of 0.7 or higher, and at least 3 fragments matched. Prior to matching, we removed library spectra whose precursor masses are more than 20 ppm away from the query spectrum. We then used an MS2 tolerance of 0.05 Da (default used^29^).
2. **Cosine similarity**^30,31^. Similar to spectral entropy, we employed a default cutoff of 0.7 or higher, at least 3 matched fragments to define confident matches, an MS1 tolerance of 20 ppm, and an MS2 tolerance of 0.1 Da.
3. **Embeddings from DreaMS**^32^. DreaMS similarities are computed by embedding mass spectra along with their MS1 precursors using the DreaMS encoder. The result is a fixed-length vector of size 1,024, which can then be queried against other embeddings using the standard cosine similarity measure widely used outside the mass spectrometry community. We applied a default DreaMS similarity cutoff of 0.9, as recommended by the authors for exact matches. DreaMS does not require any additional parameters.

We compared all three methods of spectral matching across a full range of cutoffs from 0 to 1, with a step size of 0.01, when computing ground-truth metrics. Due to the scale of the datasets we analyzed and the need to run spectral-matching algorithms multiple times throughout our study, we implemented custom spectral entropy and cosine kernels that apply an MS1 filter of 20 ppm to the spectral library before performing MS2-level matching. The kernels remain, algorithmically, equivalent to their reference implementations in the packages where they were originally implemented.

To ensure the MS1 filter does not bias results, we evaluate varying MS1 tolerances and record the corresponding maximum F1 score calculated for the ReFRAME library dataset (**Supplementary Figure 1**). We use an MS2 tolerance of 0.05 Da for spectral entropy and an MS2 tolerance of 0.1 Da for cosine similarity matching, and we provide a similar evaluation of the effect of varying this tolerance to ensure consistency in our results (**Supplementary Figure 2**). We did not implement any optimizations for computing DreaMS similarities, as after embedding the query and reference spectra with the DreaMS model, DreaMS similarities can be computed quickly using fast cosine similarity algorithms for fixed-length vectors^33^.

### Evaluation metrics

To evaluate the tools, we developed a set of metrics that capture various aspects of the feature tables generated by the tools **(Figure 1)**. Across all datasets, we report basic statistics, such as the numbers of MS1 and MS2 spectra, as well as two novel metrics we developed as proxies for MS2 quality and feature splitting (details in **Supplementary Texts 8 and 9**). In addition, we focus on MS2 characteristics (e.g., peak counts) and evaluate spectral quality through a manual expert assessment **(Supplementary File 1**). For the ground-truth datasets, we further compute precision, recall, and F1-score, comparing the spiked-in compounds, known to be present in the samples, with the set of confidently annotated structures. To avoid relying solely on the default similarity threshold for defining a confident annotation (e.g., > 0.7 spectral entropy similarity), we explored how these metrics varied over the full range of spectral similarity scores. Furthermore, we examined how their robustness changed using the three spectral-matching approaches described above to circumvent potential biases introduced by any single annotation method.

Lastly, we assessed the quantification ability and feature detection performance at varying metabolite concentrations using data collected from our two internally generated ground-truth datasets, the spiked plasma dataset and the plant natural products dataset. Additionally, to assess the ability of IM-derived CCS values to resolve co-eluting isomers in untargeted metabolomics, we performed a pairwise discrimination analysis on the same dataset. Here, we focused on structural isomers, defined as pairs with different 2D InChIKeys (i.e., connectivity) but the same formulae, corresponding to constitutional or positional isomers with the same molecular formula. For each pair, we calculated a relative ΔCCS%. The ΔCCS% was then assigned to one of four categories following Met4DX^34^: ≥4% (baseline ion mobility separation), 2-4% (partial separation), 1-2% (requiring combined LC×IM), and <1% (CCS alone insufficient, relying on chromatographic resolution). This stratification allowed us to ask whether ion mobility adds discriminatory power beyond what *m/z* (MS1) and RT alone can provide and whether that power differs meaningfully between structural isomers. It is important to note that, because the ReFRAME library ground-truth metadata did not include stereochemistry information, we did not perform a corresponding discrimination analysis for stereoisomers.

For the spiked plasma dataset, we evaluate maximum compound recall across the four different compound concentrations to determine whether the relative feature-detection performance of the tools we evaluate varies with signal intensity. To increase sample complexity, spiked-in compounds within a single set of technical replicates are not all at the same concentration; that is, a single spiked sample may contain a fraction of compounds at low concentration and another fraction at high concentration. To account for this, we compute recall at a given concentration by taking the union of the compounds spiked in at that concentration across all samples. For example, to compute recall at low concentration, we take the subset of ground-truth compounds present at low concentration in each sample, take the union across all samples, and treat the resulting set’s cardinality as the number of ground-truth compounds detected at low concentration. This enables us to compute an individual recall for each spiked-in compound concentration while maintaining the task difficulty and realism introduced by mixing spiked-in compounds of different concentrations within a single sample, rather than relying on samples with uniform concentrations of spiked-in metabolites, which would present a simpler, less realistic task.

To facilitate a rigorous assessment of quantification performance while mitigating the confounding influence of matrix effects, we evaluated the linearity of feature detection across the spiked plasma and plant natural products datasets. For the ground-truth compounds detected in the spiked plasma dataset, we calculated the coefficient of determination (R^2^) between observed feature intensities and the known on column concentrations injected. Similarly, for the plant natural products dataset, R^2^ values were computed using the three on column amounts (50, 100, and 200 ng). Under ideal conditions, a robust linear relationship is expected between these variables; therefore, we compared the distributions of R2 values across samples for each tool to gauge quantitative accuracy. Finally, we assessed performance using raw feature intensities and centered log-ratio (CLR) and robust centered log-ratio (rCLR) transformations to ensure our evaluation remained agnostic to post-processing methodologies commonly used in metabolomics research^35^.

### Data availability and implementation

We implemented the scripts and notebook in Python version 3.10. We leveraged open-source libraries, including NumPy^36^, pandas^37^, and SciPy^38^. We employed seaborn^39^ and Matplotlib^40^ for visualization. The codebase and scripts necessary to reproduce our work are available at https://github.com/enveda/tims-bench. The raw data files we processed with each of the benchmarked tools are available at the GNPS MassIVE repositories (Spiked-in NIST SRM 1950 human blood plasma <u>MSV000101615</u> and plant natural products dataset <u>MSV000101619</u>).

## RESULTS AND DISCUSSION

### Benchmarking across ten public timsTOF datasets reveals tool-dependent divergence in feature detection and MS2-based annotation

Before our work, efforts to benchmark software tools for processing untargeted metabolomics data were extremely limited. At best, among the 28 published studies in this bucket, they are split evenly between benchmarking papers and studies that develop and introduce a new tool and benchmark it against established prior tools (**Supplementary File 1**). The studies in this latter category are typically biased, since new tool papers tend to report favorable results when the investigators have free rein to optimize their own tools, unlike the other tools they are comparing. Among these tools, xcms^10^, MZmine^7^, and MS-DIAL^8^ were tested most frequently across a wide range of conventional LC-MS and GC-MS instrument heterogeneity, presenting confounders in mass accuracy, spectral acquisition rates, dynamic ranges, and sample diversity, though disproportionately so for human plasma/serum samples. However, all these benchmarking studies were limited to non-TIMS LC-MS/MS feature-detection benchmarking, except one study^9^. This clear gap in benchmarking efforts in CCS-aware feature detection, i.e., 4D peak picking (RT x *m/z* x intensity x mobility) and deconvolution relevant to TIMS-type datasets, inspired this study.

To better understand general trends across the three tools (i.e., MZmine 4.9, MS-DIAL 5.5, and MetaboScape 14.0.3), we compared them across a panel of ten public TIMS-TOF datasets **(Supplementary Table 1)**. Firstly, we investigated the total number of features at the MS1 level *(MS1 Count)* and features with MS2s *(MS2 Count)*, as well as the total number of confidently annotated features *(Annotated Count)* (**Figure 2A**).

**Figure 2.**
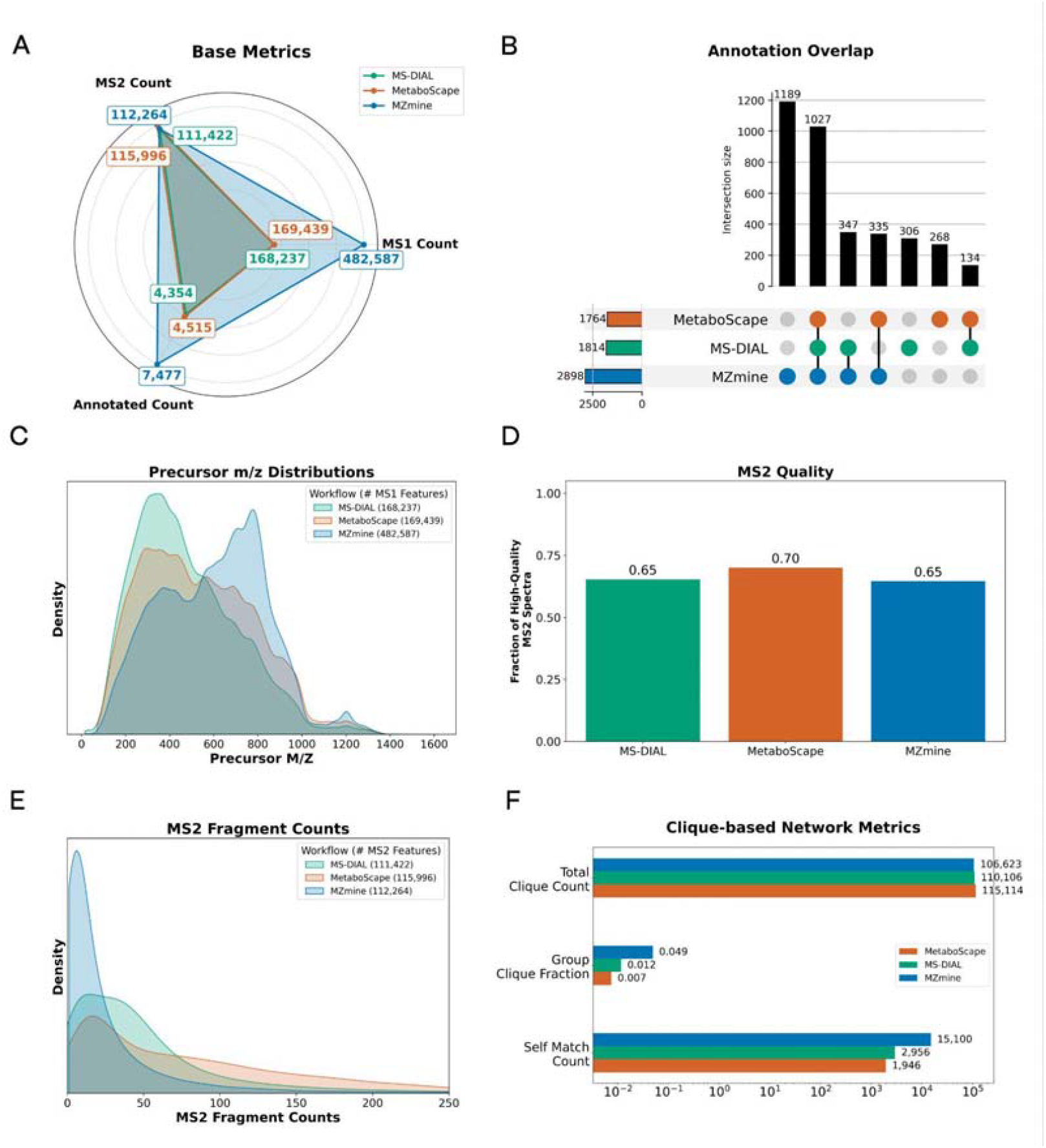
Comparative performance of metabolomics data processing tools across 10 public datasets. **A)** Aggregated counts of MS1 features (MS1 Count), MS2 feature counts (MS2 Count), and confidently annotated features (Annotated Count) across 10 public datasets. **B)** UpSet plot illustrating the overlap of unique molecular structures (i.e., InChIKey-14 identifiers) annotated for the resulting feature tables from each tool. Individual UpSet plots for each dataset are provided in **Supplementary Figure 4. C)** Precursor ion *m/z* distributions for features generated by each tool. **D)** Proportion of high-quality MS2 spectra per tool, i.e., quality as defined in the Methods. **E)** Distribution of MS2 fragments per spectrum for each tool. **F)** Clique-based network metrics summarizing spectral redundancy and consistency, including self-match count, group clique fraction, and total clique count (refer to **Supplementary Text 8** for methodological details).

Starting with the total number of MS1-only features, MZmine consistently reported the highest numbers across all ten datasets **(Supplementary Figure 3)**, followed by MS-DIAL and MetaboScape. In certain public datasets, MZmine detected up to an order of magnitude more MS1 features than the other two tools. When aggregated across all datasets, however, the MS1 feature counts were 482,587 for MZmine, 169,439 for MetaboScape, and 168,237 for MS-DIAL, a difference of approximately 3-fold between MZmine and the other two tools. While evaluating the number of MS2 features, the total counts are quite comparable across tools. Interestingly, the majority of MS1 features (MS1 counts) in MS-DIAL and MetaboScape include MS2 information, whereas in MZmine, only approximately one in four MS1 features include MS2 information which implies that MZmine reports a much greater number of MS1 features that the instrument does not acquire MS2 information for, compared to MS-DIAL and MetaboScape. Furthermore, despite the comparable MS2 feature counts, MZmine consistently yields the highest number of confident annotations (spectral entropy >= 0.7), indicating a higher annotation rate than the other two tools. However, this large gap in the number of annotated MS2 features is diminished when examining the unique number of structures being annotated (defined by InChIKey-14; **Figure 2B)**. Under these metrics, MZmine confidently annotated 2,898 unique structures (including 1,189 unique annotations that the other two tools did not annotate) while MetaboScape and MS-DIAL annotated 1,764 and 1,814 unique structures, respectively. This trend was consistent across datasets, irrespective of sample origin or chromatographic method. Interestingly, even though MZmine reported the highest number of unique annotations overall across several datasets, MS-DIAL or MetaboScape captured features that resulted in uniquely annotated structures that other tools completely missed **(Supplementary Figure 4)**. This indicates that the additional features called by MZmine may be, low-abundance (noisy), features of poor spectral quality, or different adducts of the same compounds. However, this allows MZmine to retain much more MS2 information compared to both MetaboScape and MS-DIAL. In summary, no single tool accurately captures the full extent of ionizable chemical space, and the relative strengths and weaknesses of the tools can vary depending on the complexity of the datasets under consideration.

We further estimated feature sparsity (missingness fraction), defined as the proportion of empty cells in the feature table, with 0% indicating that every feature was detected in every sample within a dataset and 100% indicating complete absence of detection. MS-DIAL and MZmine produced feature tables with comparable sparsity profiles, with medians of ~5-6%, while individual datasets reached ~17-19% for MS-DIAL and MZmine. In contrast, MetaboScape generated denser feature tables consistently, with a median missingness below 1% and no dataset exceeding ~2% (**Supplementary Figure 5**).

Next, we evaluated metrics such as precursor *m/z* distribution, MS2 quality, and MS2 fragment distribution. We observed that the three tools exhibit a comparable *m/z* distribution, with MZmine capturing more precursor m/zs between 600-1000 Da than the other two tools **(Figure 2C)**. Assessment of the MS2 quality (**see details in Methods and Supplementary Text 9)** revealed similar proportions of high-quality spectra across all tools, ranging from 65% (in MS-DIAL and MZmine) to 70% (in MetaboScape) **(Figure 2D)**. In contrast, the MS2 fragment count distribution showed that spectra generated by MZmine contained significantly fewer fragments compared to those from the other tools. This difference is likely attributable to MZmine’s MS2 merging and filtering steps, which may simplify spectra and facilitate annotation, potentially contributing to its higher annotation rates **(Figure 2E)**. Finally, we examined kernel density distributions of MS1 precursor *m/z*, retention time, MS2-selected precursor *m/z*, and CCS across the tools for each individual dataset (**Supplementary Figure 6**). Within each dataset, MS1 *m/z* and RT distributions were broadly consistent across tools, indicating that feature detection covers comparable chemical and chromatographic space regardless of the processing software. The CCS dimension also exhibited similar distributions across all three tools. This consistency confirms that ion mobility information is reliably preserved during data processing across these three software packages.

Lastly, we quantified feature splitting by a proposed clique-based network metric that measures self-match counts (defined as MS2 matches between features within narrow retention time and precursor *m/z* windows), group clique fraction (a proxy for the percentage of real features that split into duplicates) and total clique counts (a proxy for the number of real features) **(Figure 2F)** (see **Section 2.4** of the Methods for details on the metrics). MZmine showed the highest clique counts, followed by MS-DIAL and MetaboScape. Although MZmine exhibited the largest number of features overall (*MS1 count*, **Figure 2A**), the higher group clique fraction and self-match count (~5-fold greater than MS-DIAL and 7.8-fold greater than MetaboScape) confirm that a large proportion of these features is due to feature splitting originating from the same set of compounds.

To summarize, in this section we evaluated three main untargeted metabolomics tools across ten publicly available timsTOF datasets spanning plant, animal, microbial, environmental, and cell line origins on a plethora of metrics ranging from number of features to number and quality of annotations and MS2s Firstly, our evaluation shows that MZmine dominated raw feature detection with the highest MS1 counts and the highest count of confident annotations; however, 1 in 4 MS1 features were linked to an MS2 and lower fragment counts per MS2 indicates potential feature splitting or extraction of electronic instrument noise from the mass spectrometer. Secondly, MetaboScape was the most conservative in in terms of the number of reported MS1 features. Because, it reported the lowest MS1 counts, MetaboScape outputted the highest proportion of higher-quality MS2 spectra and the least feature redundancy, as measured by clique metrics. Lastly, MS-DIAL results were more of a trade-off compared to the former two tools: the majority of its reported MS1 features were linked to MS2, demonstrated the second-best feature splitting after MetaboScape (lowest group clique fraction and lowest self-match count), captured unique chemical space in annotations that the other tools missed, yet the MS2 quality fraction was the lowest.

### Ground-truth evaluation on the ReFRAME drug library demonstrates differences in true-positive recovery but preserved CCS measurements across tools

We further evaluated the three tools against three ground-truth datasets: i) ReFRAME drug library (4,723 pooled drugs), ii) spiked NIST SRM 1950 human plasma (120 metabolites at 4 concentrations in plasma as a complex matrix), and iii) a plant natural products dataset (45 natural products at 50/100/200 ng on-column in a clean background). Firstly, we evaluated the three tools on the ReFRAME library ground-truth dataset (MSV000098263), which pooled 4,723 drugs and drug-like compounds across 753 samples. Firstly, we examined the numbers of MS1 and MS2 features and annotated features **(Supplementary Figure 7)** and confirmed that the trends in this dataset are consistent with those observed in the 10 public datasets **(Section 3.1)**. MZmine reported the highest total number of MS1 and annotated features, while MS-DIAL returned the highest MS2 count among the other two tools. Next, we compared the precision-recall performance across tools as a function of spectral entropy (SE) thresholds (**Figure 3A**). The F1 curves show that MetaboScape and MZmine performed comparably, with both exceeding an F1 score of 0.8 at the 0.7 SE threshold. MS-DIAL, in comparison, presents lower F1 scores across all thresholds. It is noteworthy that at the default 0.7 SE threshold, the three tools report markedly different absolute false-positive (FP) counts (1,208 for MZmine, 861 for MetaboScape, and 586 for MS-DIAL). However, when compared with their respective true positives (TPs), the FP-to-TP ratios are similar (0.25, 0.19, and 0.21, respectively), indicating that these differences are largely proportional to the total number of annotated features produced by each tool. Importantly, TP counts (MZmine: 4,887; MetaboScape: 4,432; MS-DIAL: 2,844) substantially exceed FP counts for all tools, resulting in F1 scores of 0.854, 0.832, and 0.647, respectively. Consequently, the relative FP burden remains tightly clustered, accounting for approximately 16–20% of total annotations across all three tools. To remove annotation-method-related bias, we further compared F1-scores using two additional methods (cosine similarity and DreaMS^32^), both of which yielded similar trends (**Figure 3B**). Finally, we compared the overlap among the TPs (spiked in compounds) annotated using spectral entropy by each tool (**Figure 3C**). Overall, there was a strong consensus (i.e., 2,421 true-positive features) across the three tools, followed by the largest number of shared annotated features between MetaboScape and MZmine (1,785 features that MS-DIAL missed), while MS-DIAL and MZmine share 363 features that MetaboScape missed. Further, MZmine has 318 unique true positives, MetaboScape has 193, and MS-DIAL has 27.

Next, we assessed the accuracy of the tool-generated CCS values of the features annotated as one of the spiked-in compounds by spectral entropy. We evaluated the percent deviation between the previously reported collision cross-section (CCS) values of the ground-truth compounds and the values of the annotated features (**Figure 3D**). Here, we observed that all three tools exhibit low CCS errors, with means around 0%, indicating good precision and negligible error. All three tools exhibited high precision, with 97% (MetaboScape), 96% (MZmine), and 95% (MS-DIAL) of CCS measurements falling within a 5% error margin. However, when applying a more stringent 1% error threshold, MetaboScape achieved the highest accuracy (82%), followed by MZmine (76%) and MS-DIAL (58%) (**Supplementary Figure 8**).

Next, we wanted to evaluate the performance of the tools in extracting IM-derived CCS values as a function of the minimum ΔCCS (%) required to call a pair of isomers as discriminated against in the ReFRAME library ground-truth dataset. Before evaluating tool-level isomer discrimination performance, we measured the intrinsic CCS separability of isomeric pairs in the ReFRAME library **(Figure 3E)**. The structural isomers showed a broad ΔCCS distribution across all four categories (see **Section 2.4** in Methods), with similar numbers in each (**Supplementary Figure 9A)**. The cumulative curves quantify this gap: at the widely used 4% threshold, approximately 30% of structural isomers are IM-discriminable (**Supplementary Figure 9B)**. For structural isomer pairs, all three tools performed very closely, converging on approximately 35% discrimination at the 4% ΔCCS threshold (**Figure 3F**).

Finally, we compared the intensity distributions of all detected features with those of the true-positive annotations, observing that the intensities of true positives were significantly higher than those of the remaining low-abundance or noisy features (i.e., false negatives) **(Supplementary Figure 10)**. Furthermore, we observed that applying a uniform intensity threshold of approximately 10,000 (10^4^) across the evaluated tools consistently optimizes the resulting F1-scores. This disparity confirms that the annotated true-positive high-abundance features correspond to the pooled drugs rather than being identified by chance.

**Figure 3.**
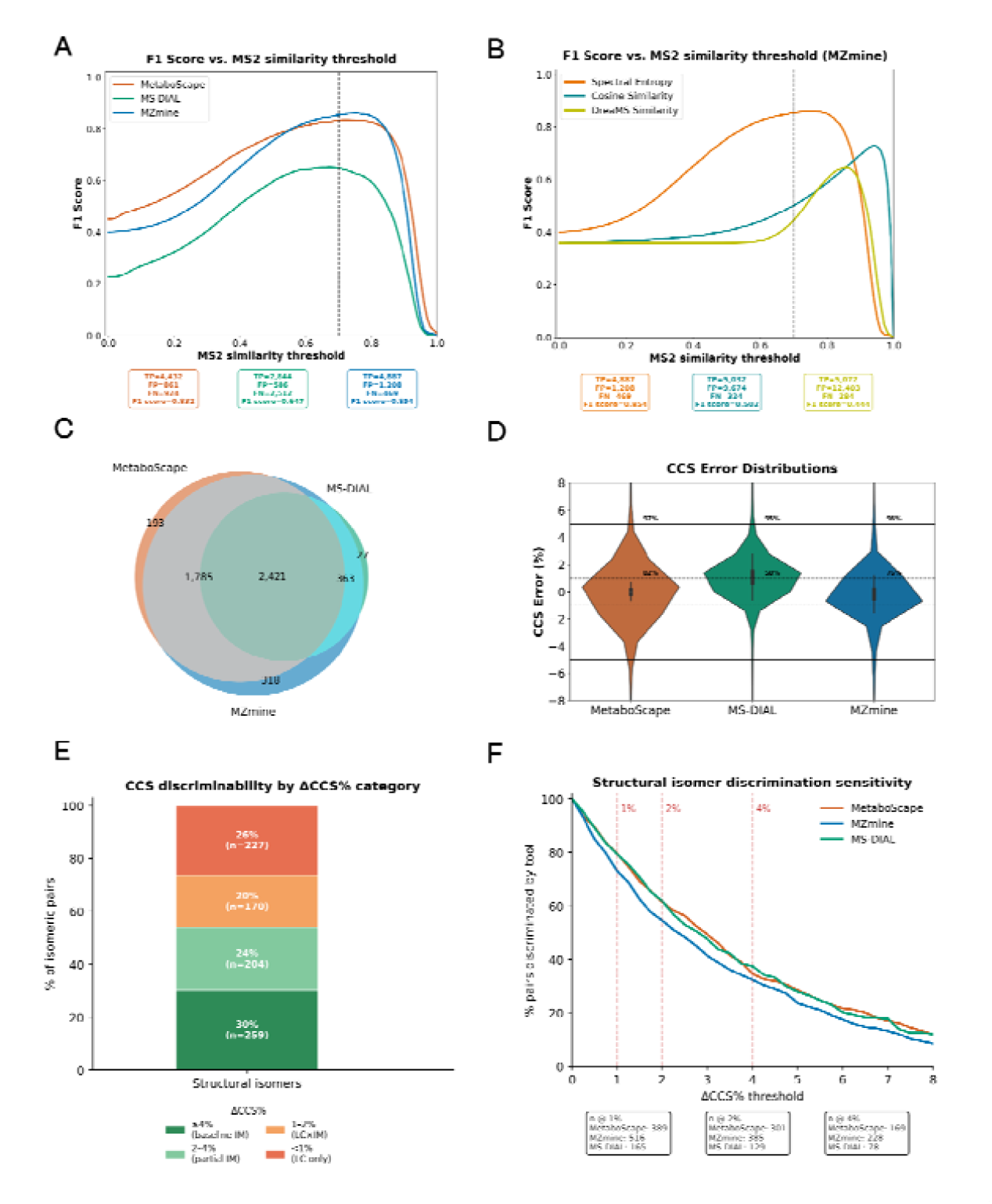
Performance evaluation and CCS accuracy on the ReFRAME library dataset. **A)** Precision-recall curves across varying similarity thresholds for each tool using spectral entropy (SE). The boxes below display the F1 scores, false positives (FP), false negatives (FN), and true positives (TP) at the default 0.7 threshold (vertical dashed line). **B)** Precision-recall threshold comparison for spectral entropy, cosine similarity, and DreaMS embeddings for MZmine. **C)** A three-way Venn diagram showing overlap of annotated true-positive features across tools using a 0.7 spectral entropy threshold. **D)** Relative CCS error (%) distribution across tools for the ReFRAME library ground-truth dataset zoomed into +-10% error. Errors at ±5 % and 1% thresholds have been highlighted, along with the percentage of data points within each threshold for each tool. **E)** Proportion of structural isomer pairs classified by pairwise ΔCCS%, binned into four categories: ≥4% (IM separation), 2-4% (partial IM separation), 1-2% (LC combined IM separation), and <1% (beyond current TIMS separation capabilities). **F)** Structural isomer discrimination performance of the tools across ΔCCS% thresholds on the ReFRAME library dataset.

### Performance on complex biological matrices and plant natural product standards remains tool-agnostic for confident annotation

Then, we evaluated the performance of the three tools against our two in-house generated datasets, the spiked-in NIST SRM 1950 human blood plasma, and the plant natural products dataset. This enabled us to rigorously test the performance of the tools in additional biological matrices, evaluate feature detection performance for compounds present in a range of concentrations in either a complex or clean background, and assess the tool’s quantification abilities.

We evaluated the feature-detection performance of each tool by calculating recall across all spike-in compound concentrations in human plasma **(Figure 4A)**. Consistent with trends observed in the ReFRAME library dataset, MS-DIAL exhibited slightly lower performance than MZmine and MetaboScape. However, the performance disparity was notably smaller than in the former dataset. Notably, the recall remained below 0.5 across the entire range of thresholds, underscoring the inherent difficulty of compound detection and annotation within a complex biological matrix such as human blood plasma.

**Figure 4.**
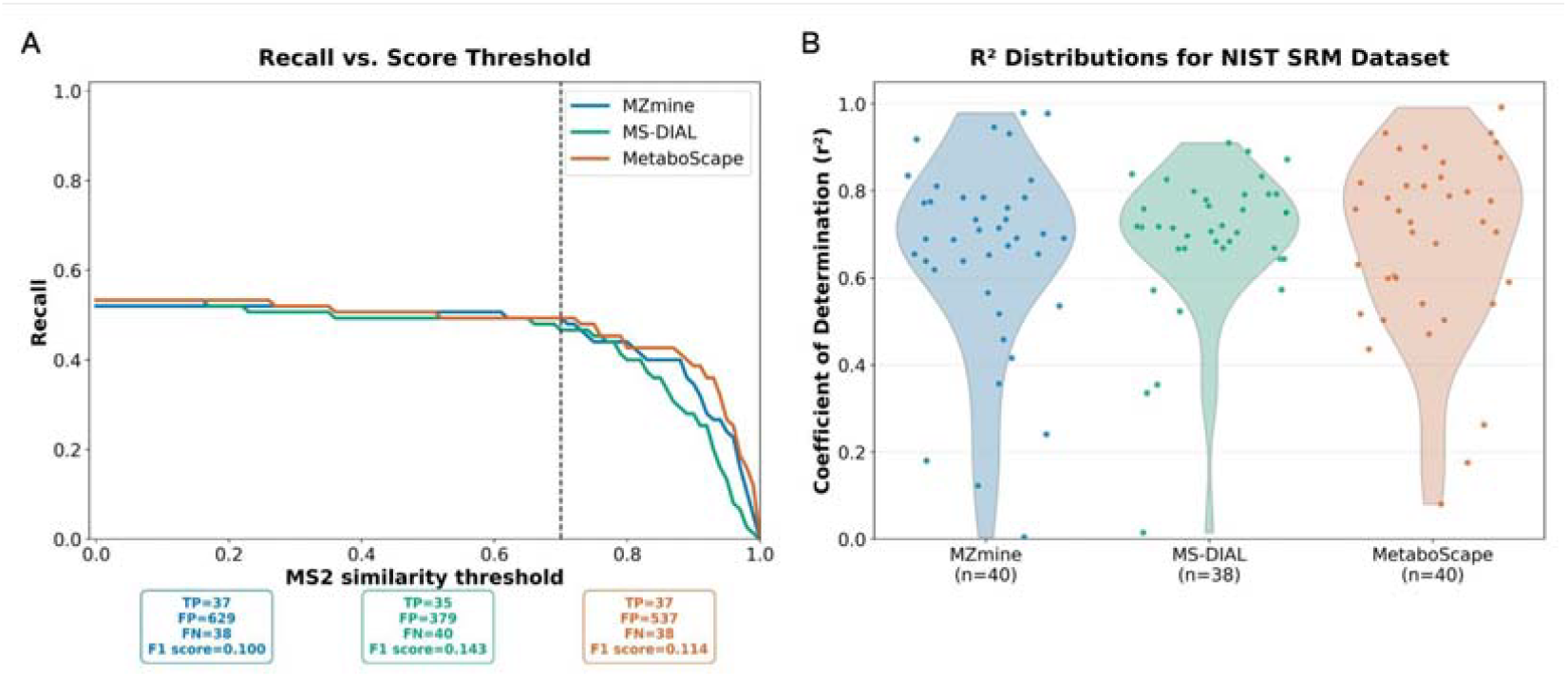
Performance evaluation and quantification accuracy on the spiked-in NIST SRM 1950 human blood plasma dataset. A) Recall curves across varying similarity thresholds for each tool using spectral entropy (SE). The boxes below display the recall, false negatives (FN), and true positives (TP) at the default 0.7 threshold (vertical dashed line). B) Distributions of the coefficient of determination (R^2^) for each tool for features annotated as ground-truth spiked compounds.

Additionally, we evaluated the correlation between the intensities of the detected ground-truth spiked-in compounds in plasma and their respective concentrations, serving as a proxy for each tool’s reporting of abundances, i.e., peak area under the curve **(Figure 4B)**. Collectively, the three tools demonstrated strong performance, with the majority of detected features (around 40 per tool) exhibiting R^2^ values above 0.6. Overall, MS-DIAL has a more compact distribution, with most values close to 0.7, whereas MZmine and MetaboScape show a wider distribution.

We did not compute precision or F1 scores for this dataset because precision requires a complete ground-truth for all compounds present, which is unavailable for the plasma background. Many spiked-in compounds naturally occur in NIST SRM 1950 human plasma (e.g., cholic acid, norepinephrine, lysine, etc.)^41^, precluding simple removal of control plasma features. Subtracting control feature intensities from spiked-in samples is confounded by sample-to-sample variation, instrument noise, and matrix effects, and any arbitrary intensity-increase threshold would differentially affect tool performance, compromising a fair evaluation.

To date, the establishment of comprehensive ground-truth or gold-standard datasets for benchmarking untargeted LC-MS/MS-scale metabolomics remains largely elusive, primarily due to the absence of standardized test materials and universally accepted metrics for rigorous evaluation^42^. While the proteomics field transitioned to robust benchmarking as early as 2016 through hybrid proteome samples of defined quantitative compositions^43^, equivalent efforts in metabolomics have been limited to inter-laboratory ring trials on reference materials such as NIST SRM 1950 human plasma. These investigations were either conducted in a targeted fashion^44^ or, more recently, via untargeted profiling^41^; however, they neither pursued ground-truth benchmarking nor utilized ion mobility-derived datasets. To address this deficiency, we generated three distinct ground-truth datasets: a spiked-in NIST SRM 1950 human plasma dataset featuring 120 metabolites within a complex biological matrix, a plant natural product mixture analyzed as pure standards to provide a zero-background control, and a large-scale drug library comprising pooled compounds to evaluate retrieval performance at scale.

To precisely assess the tool’s quantification ability in the absence of background features or sample complexity, we then computed feature-detection performance across a range of diverse plant natural products. Evaluation of feature-detection performance for each tool by calculating recall across all compound amounts showed that MZmine achieved the highest recall, closely followed by MetaboScape, where MS-DIAL has the best precision-recall balance (F1) by a wide margin **(Figure 5A)**. In addition, MZmine’s FP count is 2.4-fold of MS-DIAL’s and yields the lowest F1 on this dataset. Overall, all three tools exhibit a recall of over 0.8 with the default spectral entropy threshold of 0.7. Increasing this threshold beyond 0.9 results in a sharp drop, underscoring that a spectral entropy score around 0.7 offers a good trade-off between recall and precision, as we observed in previous datasets.

Similar to the previous dataset, we evaluated the dose-responsive increases of features (19 of the total 45 quantified natural products across three tools) from 50, 100, to 200 ng on column **(Figure 5B)**. MZmine exhibited the most favorable distribution of R^2^ values, identifying only a single feature below the 0.4 threshold. In contrast, MetaboScape and MS-DIAL each reported several low-correlation outliers. Notably, MZmine also captured the largest subset of features with R^2^ values exceeding 0.9. MetaboScape maintained a robust distribution, with the majority of features showing strong quantitative correlation, whereas MS-DIAL exhibited a more dispersed performance profile. These findings underscore that, despite robust feature detection sensitivity, maintaining quantitative consistency across varying metabolite concentrations remains a significant analytical challenge.

**Figure 5.**
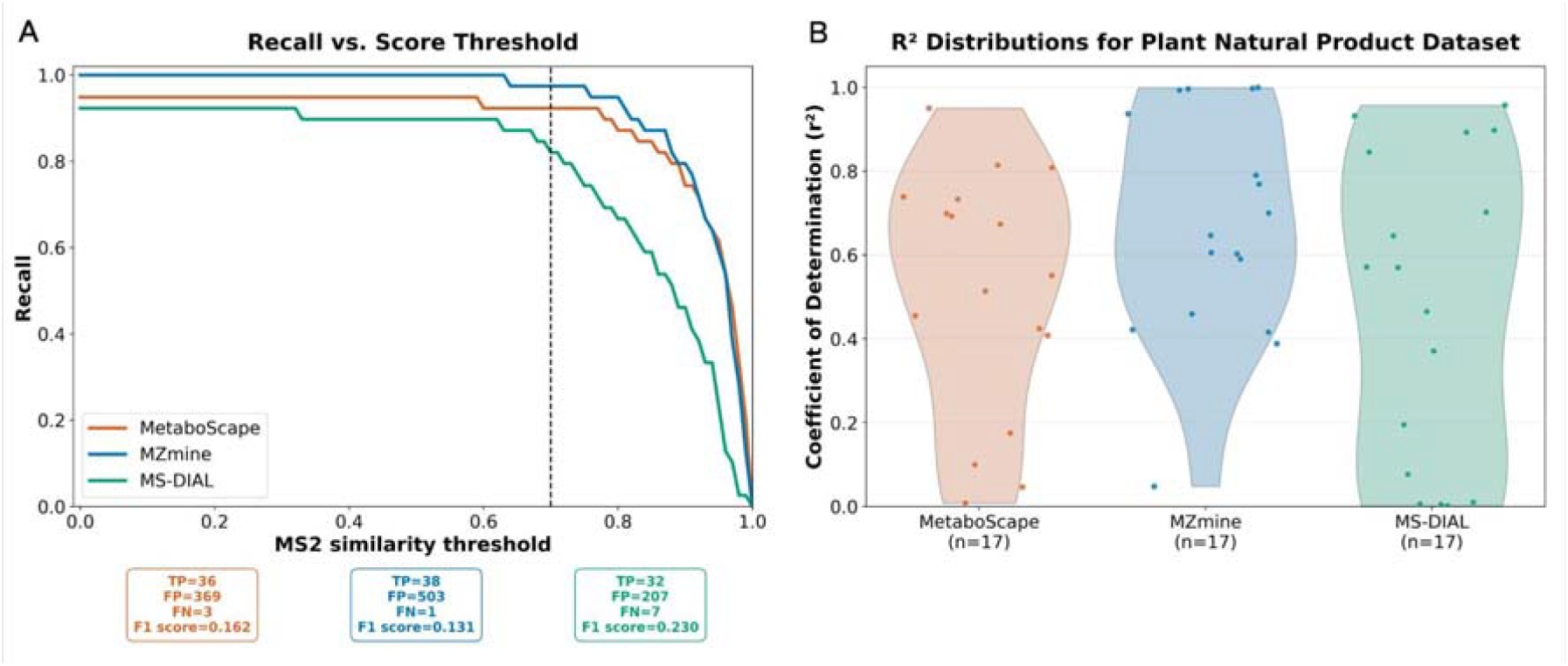
Performance evaluation and quantification accuracy on the plant natural products dataset. A) Recall curves across varying similarity thresholds for each tool using spectral entropy (SE). The boxes below display the recall, false negatives (FN), and true positives (TP) at the default 0.7 threshold (vertical dashed line). B) Distributions of the coefficient of determination (R^2^) for each tool for features annotated as ground-truth spiked compounds. For each unique compound, the adduct with the highest total spectral entropy score across all samples was chosen, with each adduct needing to meet a minimum threshold of 0.7 SE score to be included.

### Implications, future directions, and study limitations

Results indicated that MZmine performed best for raw detection sensitivity on the ReFRAME dataset and achieved the highest recall on the plant natural products dataset; however, on the latter, it also exhibited high variability in quantification, with a wide spread of R^2^ values across spiked compounds. MetaboScape matched MZmine’s F1 score on the ReFRAME and showed better CCS accuracy than the other tools, but exhibited slightly lower R^2^ distributions for dose-response linearity in the plant natural products dataset. Finally, MS-DIAL showed consistent yet lower performance on detection metrics, with lower F1 scores on the ReFRAME dataset across all spectral matching thresholds, even though it produced the highest MS2 counts **(Supplementary Figure 7)**. On quantification, it ranked closer to the other two tools, and its strengths were in MS2 acquisition rather than precision in feature calling or annotation.

Despite being time-consuming and impractical at scale due to compute requirements, the use of multiple untargeted LC-MS/MS data processing tools has become increasingly common in natural products and phytochemistry research. For instance, a recent effort that led to the isolation of cytotoxic daphne diterpenoids from *Daphne altaica* relied on processing from both MZmine and MS-DIAL^45^. Another effort on DIA-MSE data, IntOpenStream used a workflow that explicitly combined MZmine 3 and MS-DIAL applied to the analysis of 60 plant extracts from the *Ocotea* genus^46^. In other plant metabolomics efforts, the investigators used MZmine to align DDA and DIA files and integrate feature lists, while MS-DIAL was used for DIA small-molecule processing^47^. Thus, the complementarity of the processing tools for extracting features that represent different chemical spaces is well known.

Additionally, we compared three spectral matching methods (cosine, spectral entropy, and DreaMS) across the tools. Bushuiev et al.^32^ previously noted DreaMS’s superiority in structural correlation and retrieval, while Li et al.^29^ demonstrated low false discovery rates using entropy-based matching for natural products. Our findings indicate that a spectral entropy similarity of ~0.7 provides the optimal precision-recall balance, allowing us to recommend thresholds based on performance on a controlled ground-truth dataset. Regarding evaluation metrics, one study found that the linear weighted moving average (LWMA) algorithm from MS-DIAL achieved the highest TP and recall rates across 10 diverse metabolomics datasets^48^. Another study on environmental non-targeted screening datasets reported recalls of 88% for Compound Discoverer, 83% for enviMass, 82% for MZmine2, and 64% for XCMS Online, with feature overlap among all four programs at ~10%. For each software, only 40% and 55% of features did not match those of any other program^49^. However, none of these efforts address the fourth dimension of IM and are outdated versions of the current software we evaluated here.

Our study has several limitations that future works should aim to address. Since the main focus of the study is to evaluate the feature-detection ability of the tools we tested, our quantification performance evaluations are relatively simple. Future work that aims to evaluate quantification ability further would benefit from collecting ground-truth data by spiking a set of metabolites of known concentration into a complex background containing a disjoint set of metabolites. This would allow for the tools that we evaluated to be tested on a more challenging quantification task. Furthermore, we did not analyze any negative mode data because the number of negative mode timsTOF data sets deposited in the public domain is small, and because spectral libraries containing negative mode MS2 spectra are relatively sparse, severely constraining our ability to annotate the tool outputs. For reference, the library we used in this study, comprising all spectra from public libraries (see Methods), contained 1,005,059 negative-mode spectra from 47,275 unique compounds. In comparison, the same library contains 2,893,050 positive mode spectra from 99,664 unique compounds. We also did not analyze data from other types of instruments capable of collecting IM data, since we only have access to timsTOF instruments and thus could not generate ground-truth data for them, and because there are relatively few publicly available datasets from other types of instruments that support mobility data. The same goes on to explain that we did not account for other MALDI- or imaging MS-associated IM datasets or data-independent acquisition (DIA)-generated metabolomics datasets due to the lack of publicly available, ground-truth datasets. Moreover, all the (MS2 spectra to InChiKey-14 matches) metabolite annotations reported in this study correspond to Metabolomics Standards Initiative (MSI) Level 2 confidence; the study did not account for matches that take into consideration retention time (RT) and missed out on comparing the CCS values when not part of a spectrum in the reference libraries. However, limiting our study to a single instrument type allows for a more rigorous evaluation across a diverse range of tasks, rather than sparse evaluations across multiple instrument types. Moreover, we inadvertently realize that all three ground truth datasets were generated using very comparable chromatography and TIMS instrumentation, thus rendering them analytically less diverse and biased toward these analytical methods. Lastly, we did not perform systematic parameter optimization for each tool-dataset combination due to computational constraints and instead adopted HRMS analysis settings standard in the field.

## CONCLUSIONS

In summary, no single data-processing tool captured the full extent of the ionizable chemical space, as demonstrated by our evaluation across 13 public and ground-truth datasets spanning a wide range of biological matrices and organismal diversity. While each tool offers unique strengths, they also present distinct limitations when processing TIMS-amenable datasets, highlighting significant scope for future refinement. We observed that precision–recall trade-offs were highly dataset-dependent; specifically, biologically complex matrices yielded lower recall, whereas cleaner backgrounds facilitated a clearer differentiation of tool performance. Given that these tools offer unique coverage of different chemical spaces, leveraging the union of their results remains informative; however, such an approach is often impractical for large-scale datasets and routine analytical tools.

## ASSOCIATED CONTENT

### SUPPORTING INFORMATION

#### SUPPLEMENTARY FIGURES

**Supplementary Figure 1**. Total number of detected MS1 features (MS1 Count), features with associated MS2 spectra (MS2 Count), and confidently annotated features (Annotated Count), aggregated across all ten public datasets.

**Supplementary Figure 2**. Overlap of unique molecular structures annotated by each workflow for each of the ten public datasets represented as an UpSet plot based on InChIKey-14 identifiers from all confidently annotated features.

**Supplementary Figure 3**. Feature sparsity calculated across all ten public datasets.

**Supplementary Figure 4**. Distributions for each dataset for MS1 (in Da), RT (in min), MS2 (in Da), and CCS (in Å) scan range.

**Supplementary Figure 5**. Total number of detected MS1 features (MS1 Count), features with associated MS2 spectra (MS2 Count), and confidently annotated features (Annotated Count) for the three groundtruth datasets.

**Supplementary Figure 6**. Distributions of the relative CCS error (%) for each tool, represented as violin plots. The total number of features annotated as ground-truth compounds is displayed at the top.

**Supplementary Figure 7**. Intrinsic CCS discriminability of isomeric pairs in the ReFRAME library ground-truth dataset.

**Supplementary Figure 8**. Intensity distribution between the detected features and the true-positive annotations (spiked-in compounds) for three ground-truth datasets (ReFRAME library, Spiked-in NIST SRM 1950 human plasma, and plant natural products) across the tools.

**Supplementary Figure 9**. Selection of optimal MS1 tolerances based on their corresponding maximum F1 score calculated for the ReFRAME library dataset.

**Supplementary Figure 10**. Selection of optimal MS2 tolerances based on their corresponding maximum F1 score calculated for the ReFRAME library dataset.

#### SUPPLEMENTARY TEXT

**Supplementary Text 1**. MS2 Quality metric.

**Supplementary Text 2**. Feature splitting proxy metrics.

**Supplementary Text 3**. National Institute of Standards and Technology (NIST) Reference Material 1950 (SRM 1950) – Metabolites in frozen human plasma.

**Supplementary Text 4**. Preparation of plant natural products.

**Supplementary Text 5**. Untargeted LC-MS/MS acquisition methods.

**Supplementary Text 6**. Parameters used to run MetaboScape.

**Supplementary Text 7**. Parameters used to run MS-DIAL.

**Supplementary Text 8**. Parameter file used to run MSDIAL.

**Supplementary Text 9**. Parameters used to run MZmine.

#### SUPPLEMENTARY TABLES

**Supplementary Table 1**. List of datasets used for the study.

**Supplementary Table 2**. List of plant-derived natural products spike-in ground-truth data with 45 compounds injected at three on column amounts (50, 100, and 200 ng).

**Supplementary Table 3**. List of adducts used for feature calling in all three software tools (MZmine, MS-DIAL, and MetaboScape).

#### SUPPLEMENTARY FILES

**Supplementary File 1**. Meta-data of 28 publications where benchmarking of software were reported with non-TIMS untargeted LC-MS/MS metabolomics datasets.

**Supplementary File 2**. List of 126 metabolites and the concentrations used, spanning six biochemical classes, spiked into NIST SRM 1950 human plasma samples.

## AUTHOR CONTRIBUTIONS

A.A. and D.D-F. conceived the project. P.R., D.D-F., and B.B.M. designed the experiments. V.D. and A.R. generated or assisted in generating the three ground-truth datasets. P.R., Y.G., and B.B.M. identified and prepared the public datasets. B.B.M. and V.D. processed the datasets in MetaboScape and MS-DIAL. P.R. processed the datasets in MZmine. P.R. analyzed the datasets and conducted the evaluation. P.R., Y.G., and D.D-F. generated the visuals. P.R. defined the evaluation metrics in collaboration with all authors. A.A., K.W, and P.C.D. provided feedback during the project. The manuscript was initially drafted by P.R., Y.G., D.D-F., and B.B.M. and edited through contributions of all authors. The manuscript was written through contributions of all authors. All authors have given approval to the final version of the manuscript.

## ACKNOWLEDGMENTS

We extend our sincere gratitude to the Bruker (MetaboScape support) team for their responsive feedback and assistance with our inquiries. We also thank Tito Damiani from IOCB Prague for his extensive feedback and comments. We thank Mason Victors for helping design the spiked plasma dataset. We also would like to thank our colleagues at Enveda for their valuable suggestions and feedback.

## ABBREVIATIONS

TIMS: trapped ion mobility spectrometry; IMS
CCS: collision cross section
IM-MS: ion mobility mass spectrometry
RT: retention time
NIST: National Institute of Standards and Technology
SRM: standard reference material

