## Supplementary Information for "TIMS-Bench: Towards community standards for benchmarking untargeted trapped ion mobility metabolomics tools and datasets"

**Supplementary File**

### **Supplementary Figures**

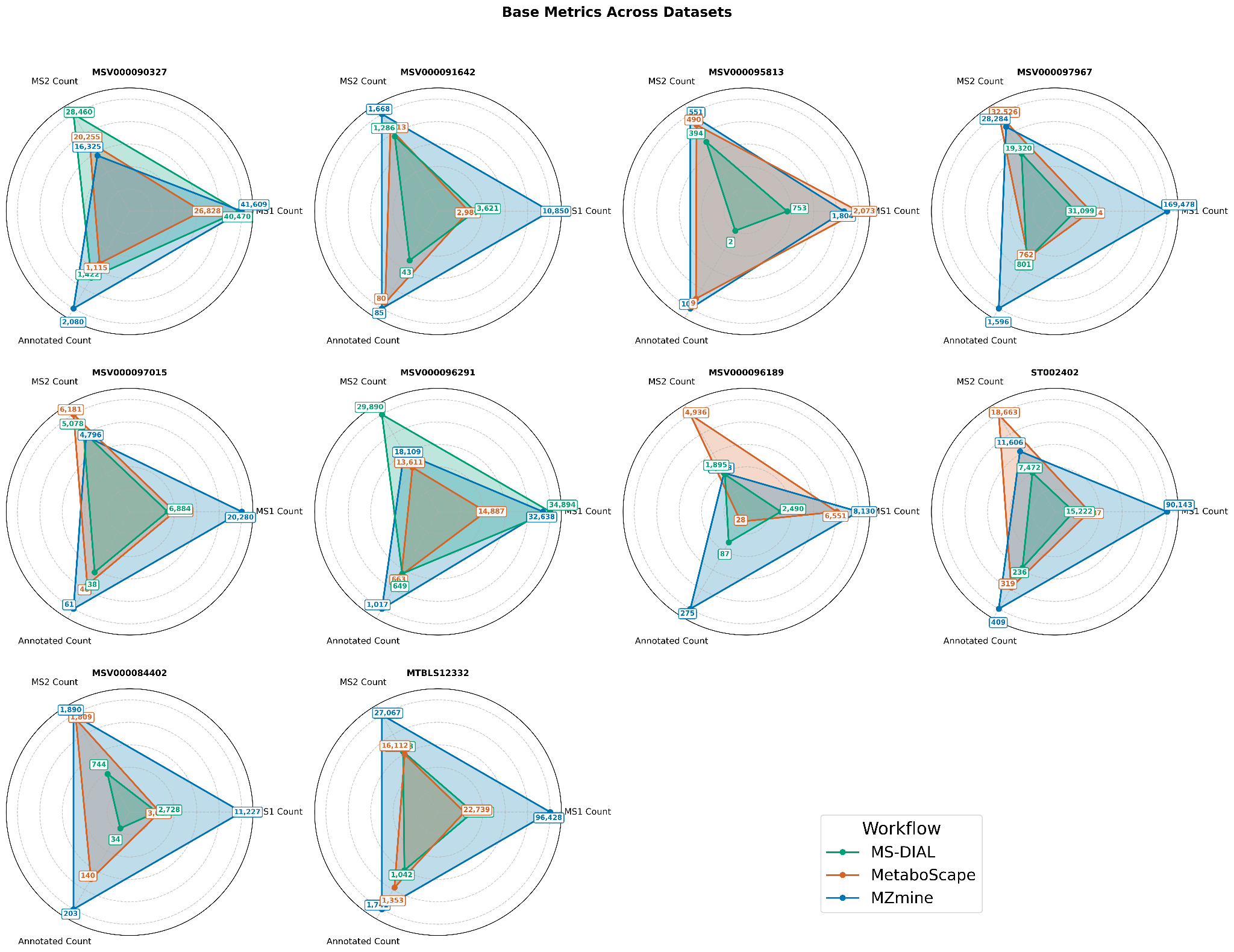

#### **Supplementary Figure 1. Total number of detected MS1 features (MS1 Count), features with associated MS2 spectra (MS2 Count), and confidently annotated features (Annotated Count), aggregated across all ten public datasets.**

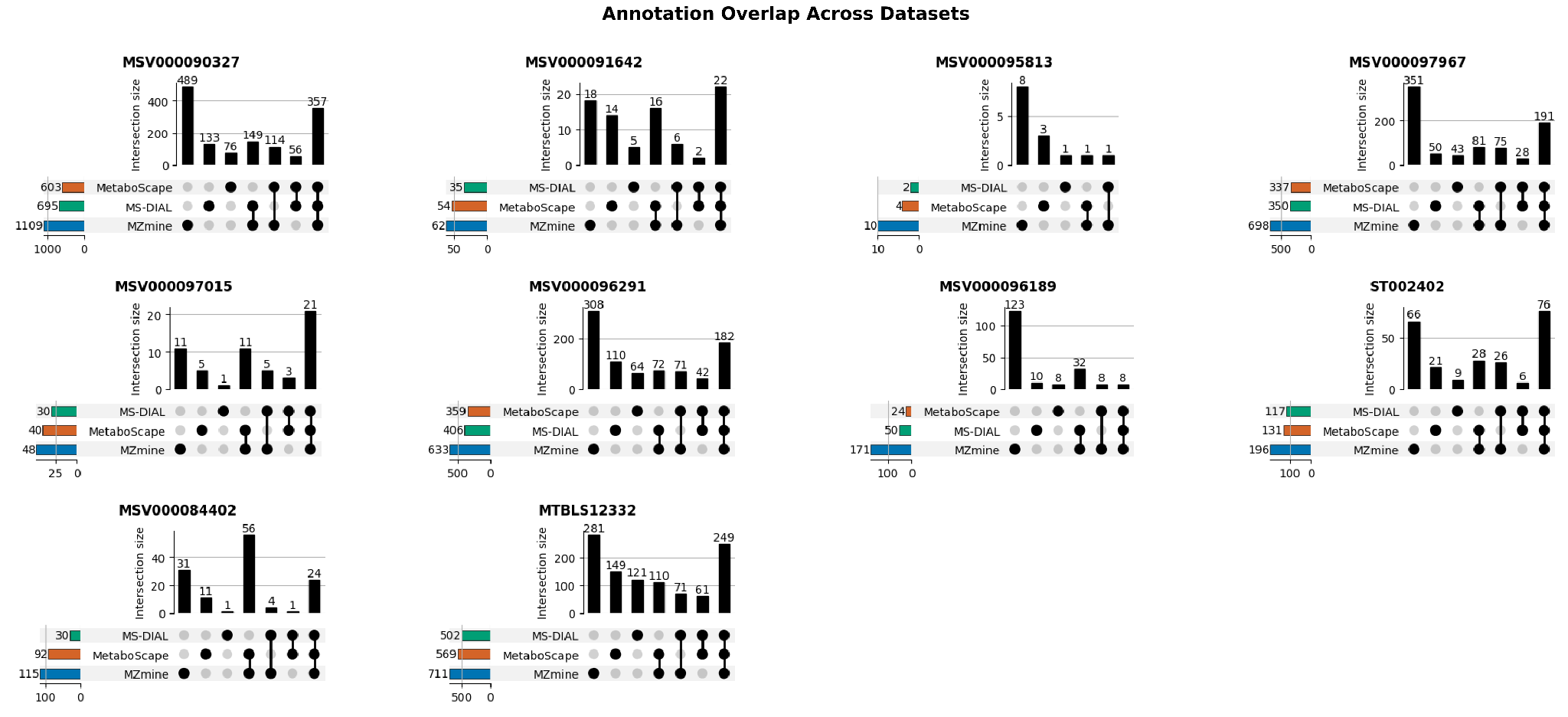
**Supplementary Figure 2. Overlap of unique molecular structures annotated by each workflow for each of the ten public datasets represented as an UpSet plot based on InChIKey-14 identifiers from all confidently annotated features.**

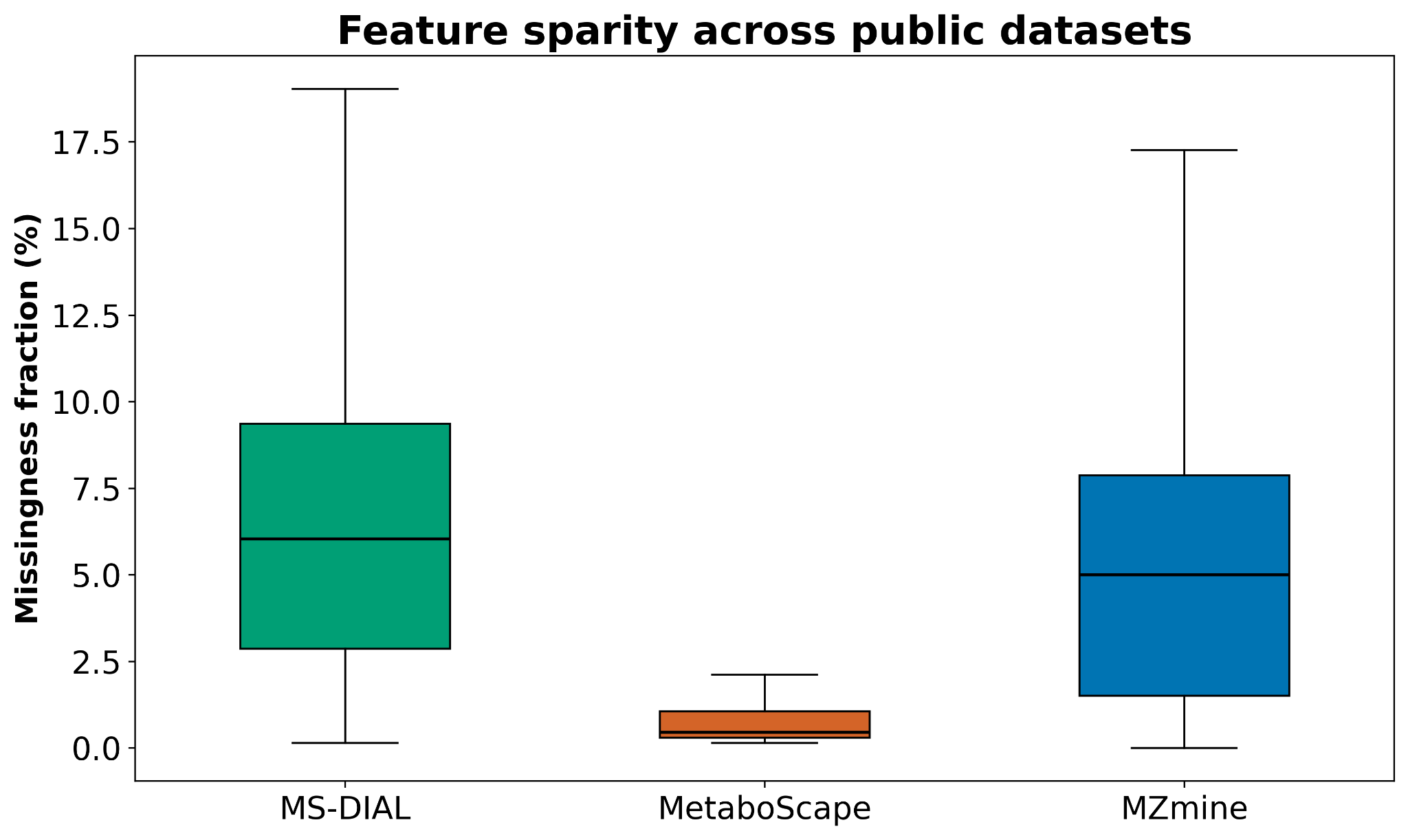

**Supplementary Figure 3. Feature sparsity calculated across all ten public datasets.**

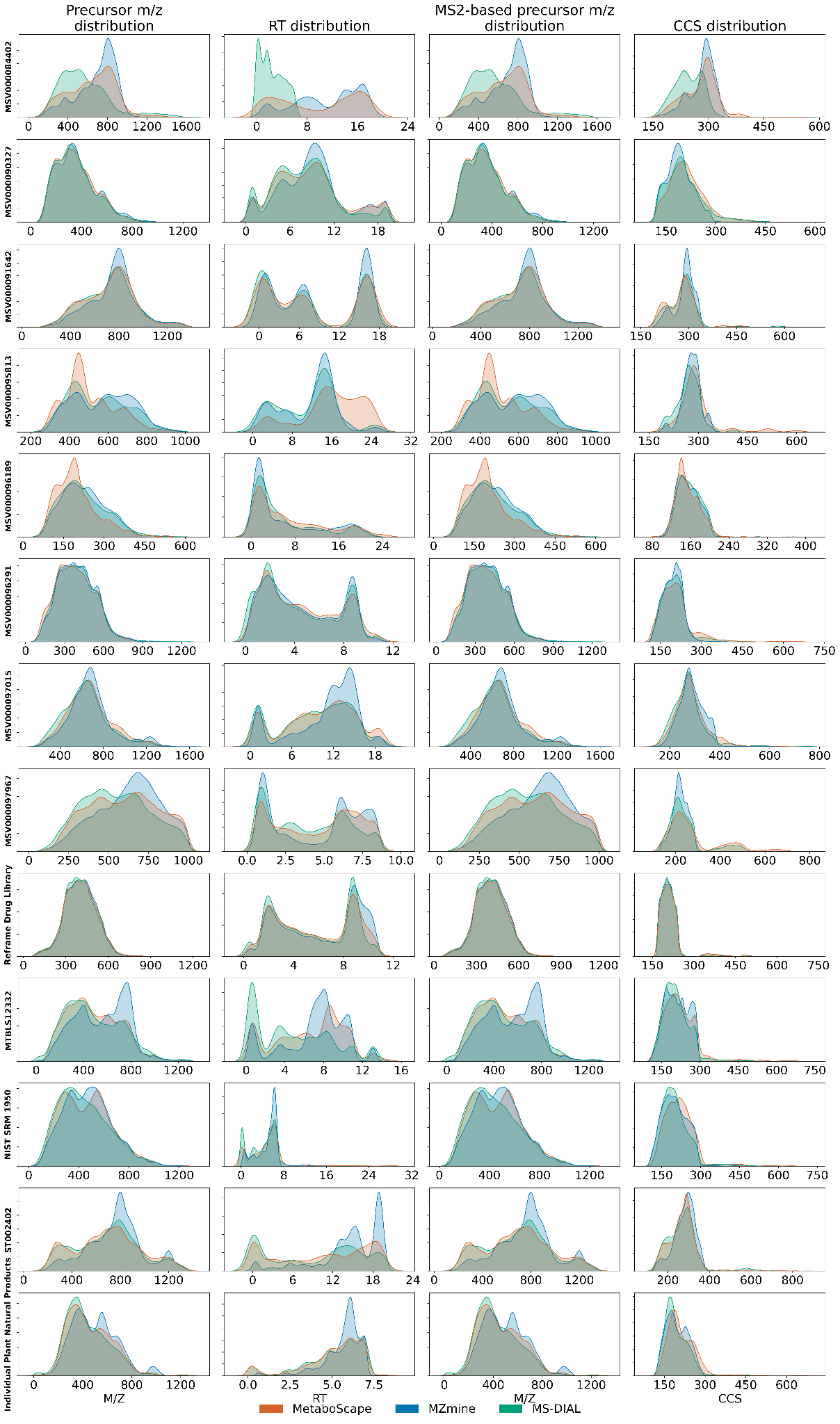

### **Supplementary Figure 4. Distributions for each dataset for MS1 (in Da), RT (in min), MS2 (in Da), and CCS (in Å) scan range.**

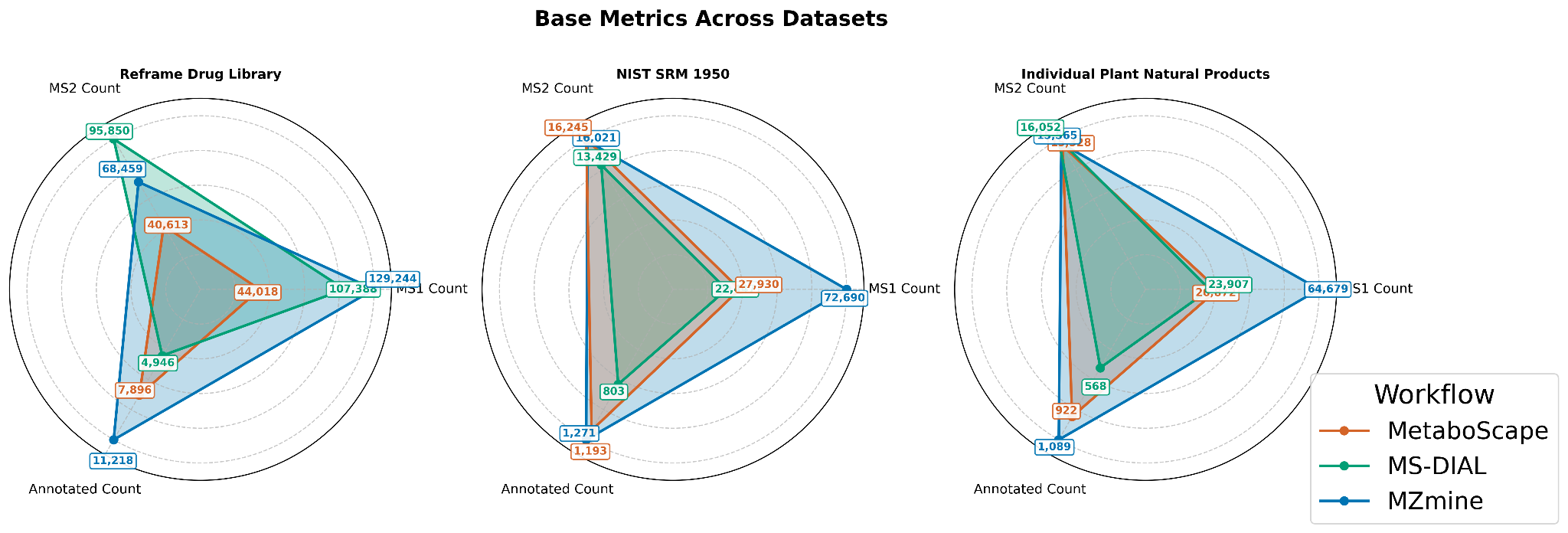

**Supplementary Figure 5. Total number of detected MS1 features (MS1 Count), features with associated MS2 spectra (MS2 Count), and confidently annotated features (Annotated Count) for the three groundtruth datasets.**

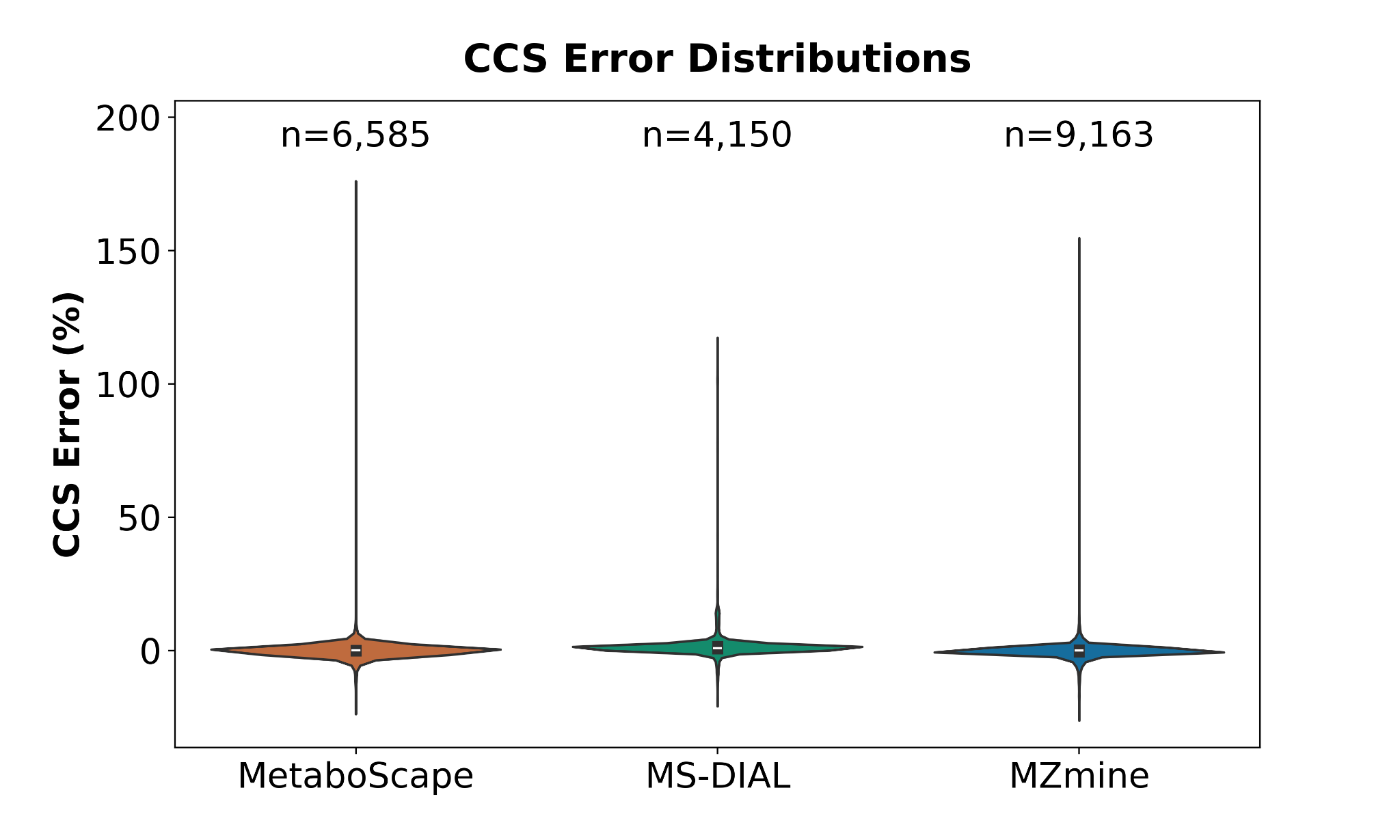

**Supplementary Figure 6. Distributions of the relative CCS error (%) for each tool, represented as violin plots. The total number of features annotated as ground-truth compounds is displayed at the top.**

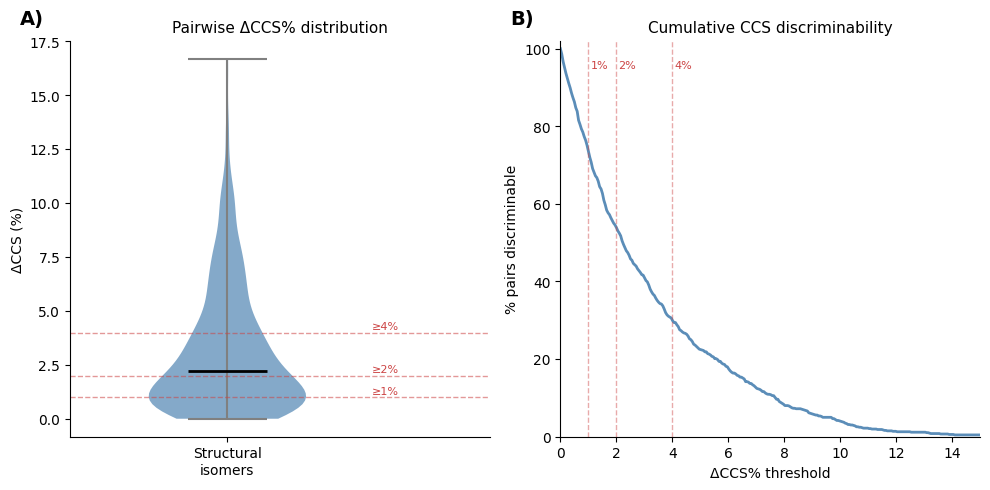

**Supplementary Figure 7. Intrinsic CCS discriminability of isomeric pairs in the ReFRAME library ground-truth dataset. A)** Violin plots of plots of pairwise ΔCCS% distributions for structural isomers; horizontal dashed lines mark the 1%, 2%, and 4% thresholds, and **B)** Cumulative discriminability curves showing the fraction of structural isomeric pairs with ΔCCS greater than or equal to a given threshold across ΔCCS% thresholds from 0 to 15%.

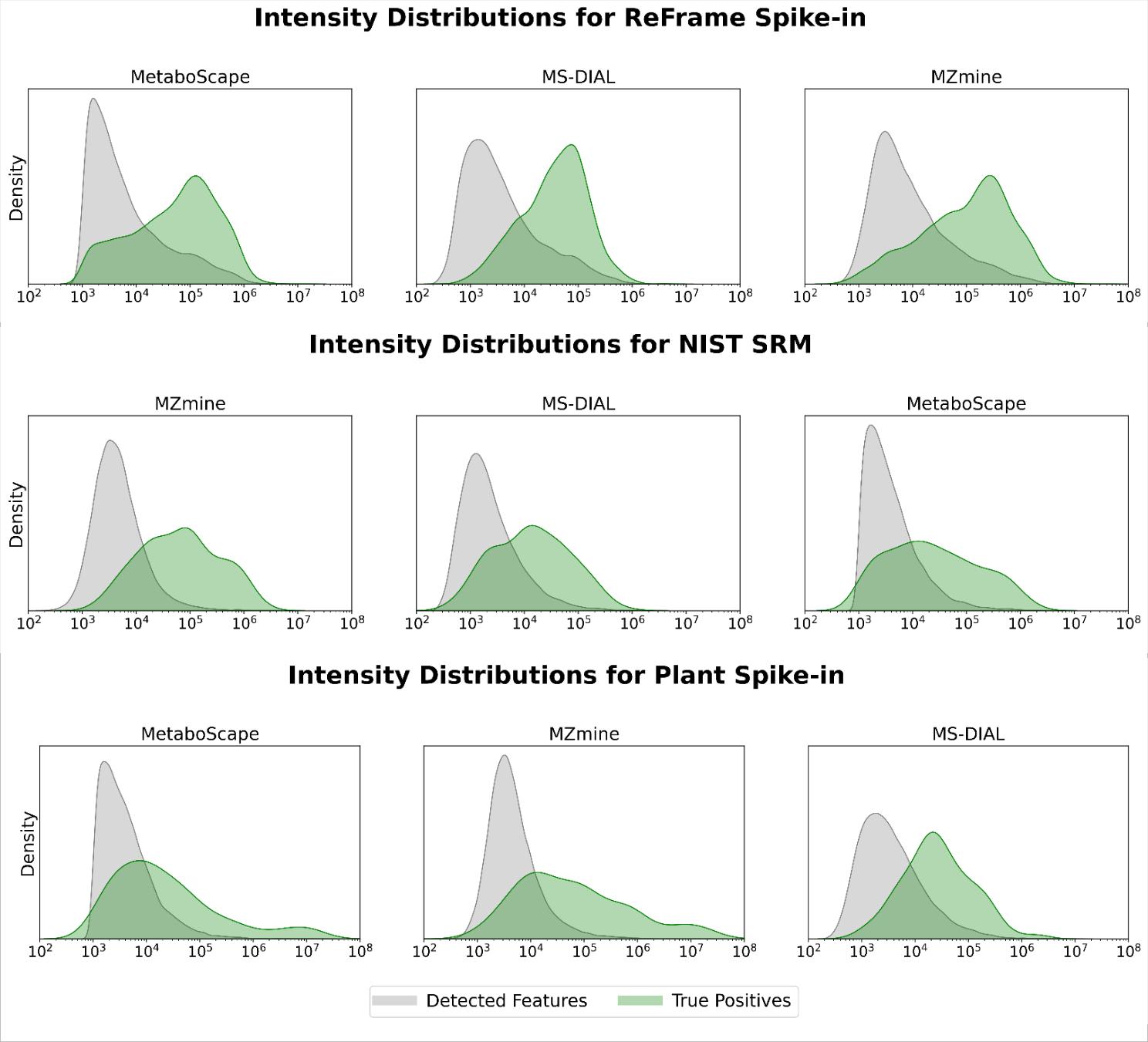

**Supplementary Figure 8. Intensity distribution between the detected features and the true-positive annotations (spiked-in compounds) for three ground-truth datasets (ReFRAME library, Spiked-in NIST SRM 1950 human plasma, and plant natural products) across the tools.** In each instance, the observed distributions exhibit sufficient separation, facilitating the visual identification of an intensity threshold that optimizes the distinction between them and maximizes the resulting F1-score. It is interesting that this threshold is around 10,000 (10^4^).

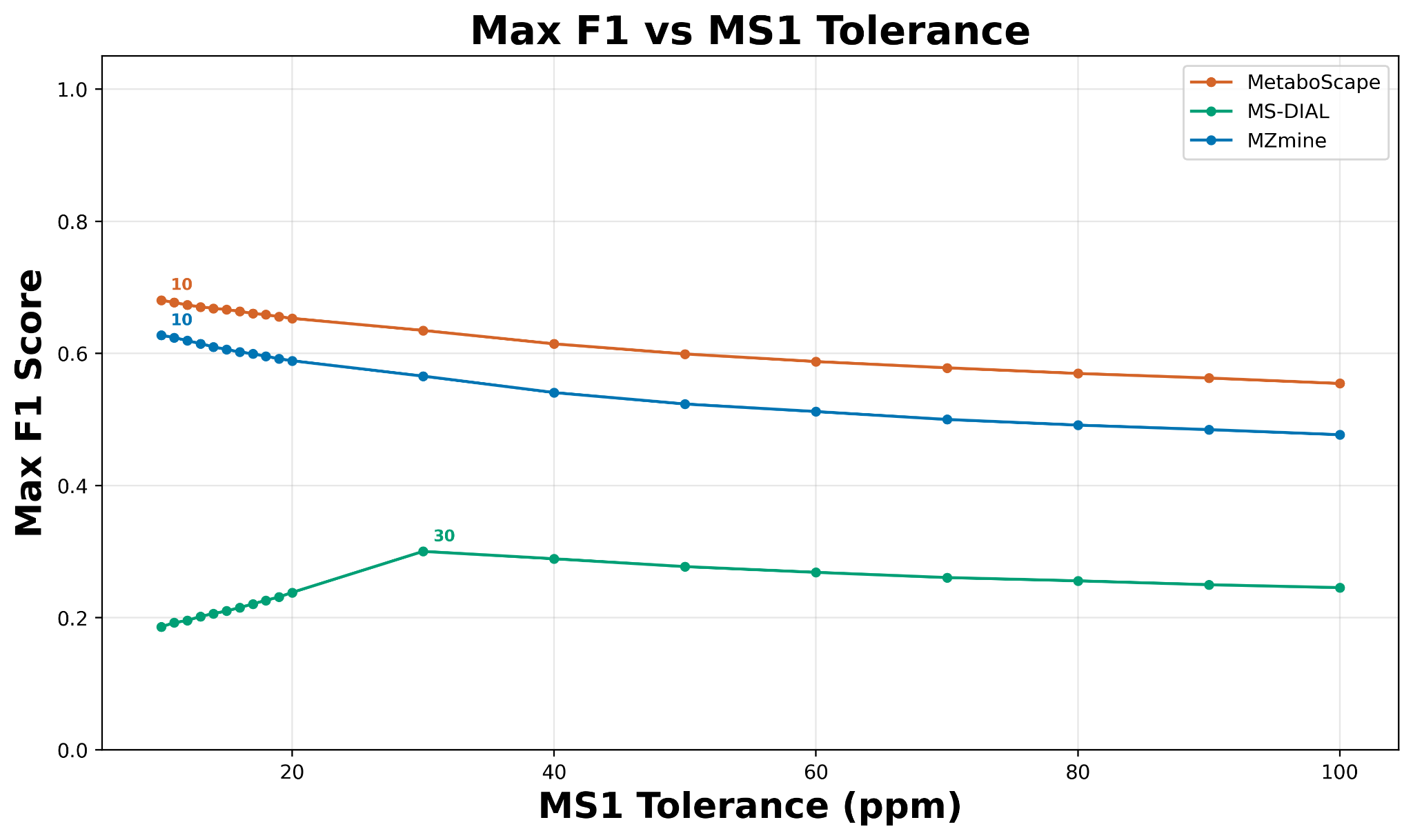

#### **Supplementary Figure 9. Selection of optimal MS1 tolerances based on their corresponding maximum F1 score calculated for the ReFRAME library dataset.**

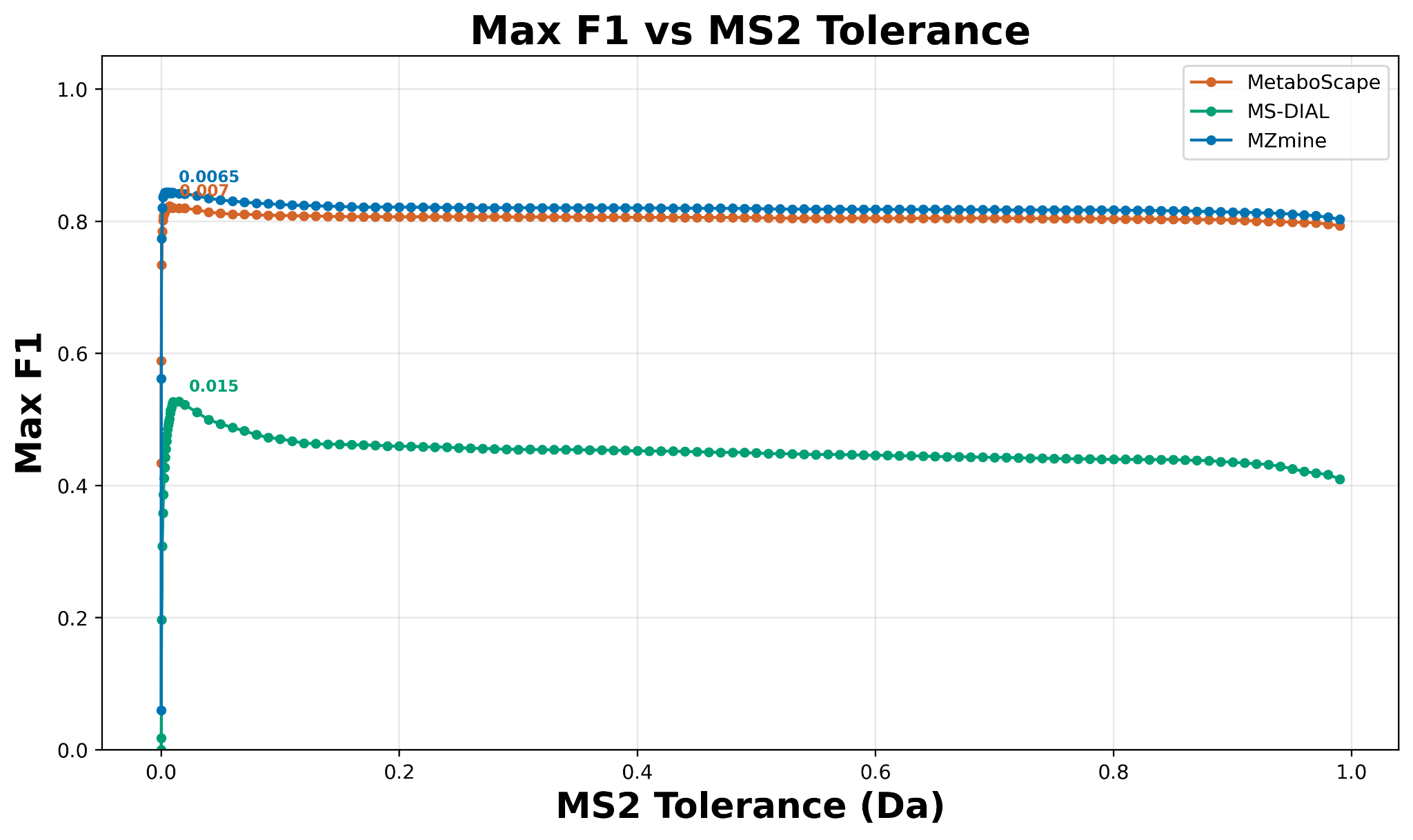

#### **Supplementary Figure 10. Selection of optimal MS2 tolerances based on their corresponding maximum F1 score calculated for the ReFRAME library dataset.**

### **Supplementary Text**

#### **Supplementary Text 1. MS2 Quality metric.**

The MS2 quality assessment follows a multi-step evaluation process designed to identify high-quality spectra suitable for reliable metabolite identification. First, spectra are evaluated using spectral entropy, where a spectral entropy higher than 1.5 is classified as “Good”, while MS2s with a lower value are classified as “Bad”. However, even spectra that pass the initial *spectral entropy* threshold undergo additional quality filters to identify problematic patterns. Thus, we also considered an MS2 as “Bad” if they exhibit uniform high peaks (where all normalized intensities exceed 50% of the maximum with at least 10 peaks), which typically indicates instrumental artifacts or contamination, or if no individual peak intensity exceeds an absolute threshold (default 200), suggesting insufficient signal for reliable fragmentation pattern analysis. This comprehensive approach ensures that only spectra with both sufficient signal strength and meaningful fragmentation patterns are classified as “Good” quality for downstream annotation and identification workflows.

**Supplementary Text 2. Feature splitting proxy metrics.**

To evaluate feature splitting, we developed a new metric called "clique cover". For a given output feature table, let N be the number of features associated with MS2s that are selected. We then construct a graph that can be represented by an N x N adjacency matrix, where edges are drawn between any two features that have (i) a spectral entropy matching score greater than 0.7, (ii) an MS1 precursor tolerance less than or equal to 20 ppm, (iii) an RT tolerance less than 0.1 minute, and (iv) a CCS tolerance less than 1%. We aim to only draw edges between two features that are highly likely to be associated with the same molecule and adduct. After constructing this graph across all MS2 features, we aim to break it apart into cliques. A clique is a subgraph in Kn, or a complete graph, meaning that every node (or feature) within a clique has an edge to every other node.

We interpret a clique of features as a set of nodes that are likely to correspond to the same molecule and adduct, since they share edges. Thus, we seek to partition an input graph of features into the smallest possible number of cliques, with the number of cliques interpreted as the number of truly unique features in a tool's output. This problem, known as the minimum clique cover problem, is NP-hard (Karp, 1972; Dau *et al.*, 2020).

To tractably solve the minimum clique cover problem, we use a greedy heuristic that provides an approximate solution serving as an upper bound. We give a formal definition of the problem and the algorithm used below.

Problem Statement: Given a graph G(V, E), a clique in the graph is defined as a set of vertices S ⊆ V such that G[S], the induced subgraph of S, forms a complete graph, that is, there exists an edge between any two vertices in G[S]. The minimum clique cover problem seeks to find the smallest number of cliques that cover the graph; that is, the smallest number of cliques such that the union of their vertex sets equals V and their vertex sets are pairwise disjoint.

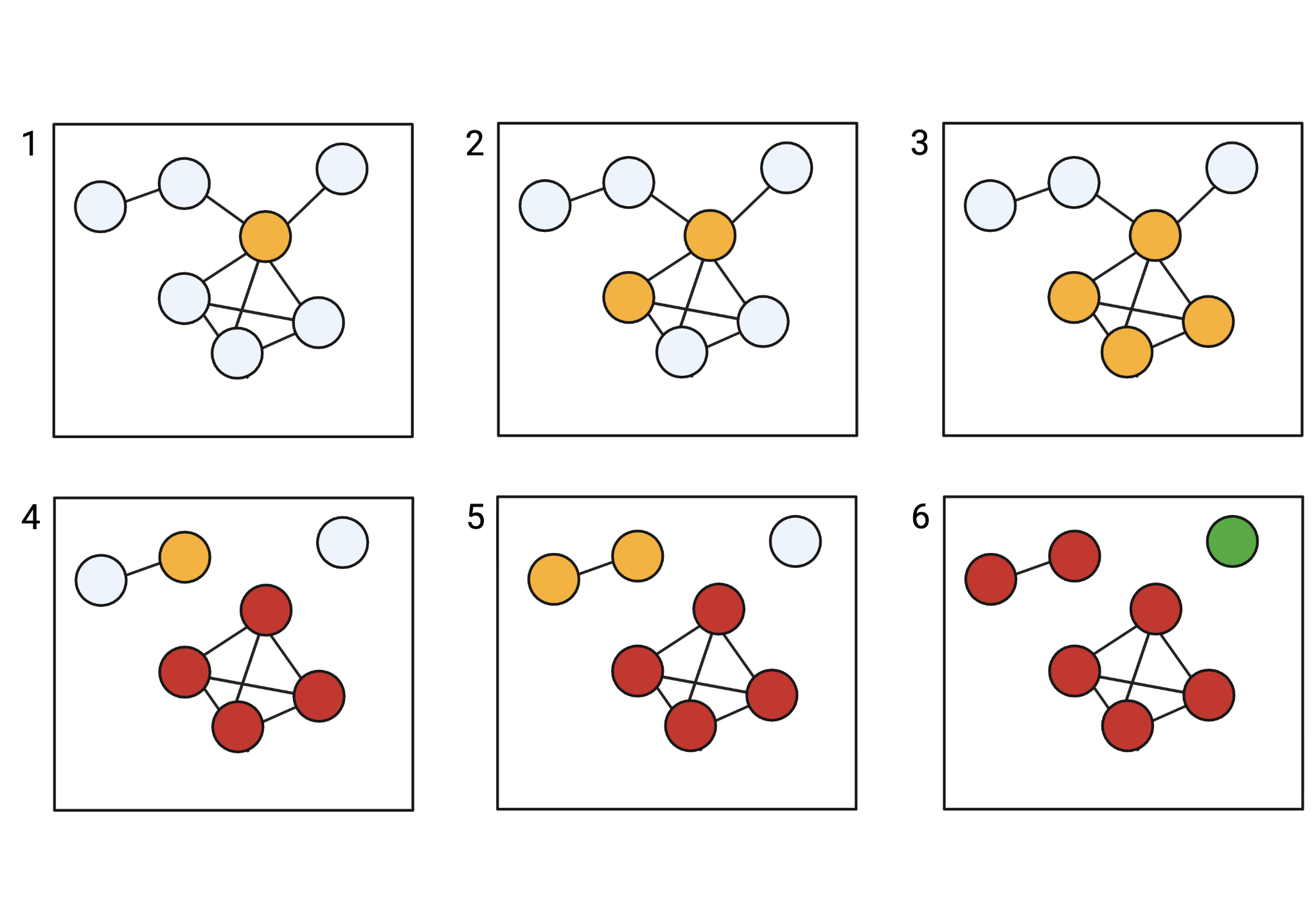

**Figure: Greedy Minimum Clique Cover Visualization.** 1. Given an input graph, the node with the greatest number of neighbors is selected and added to the current clique. 2. Next, within the remaining unprocessed nodes, the node with the greatest number of neighbors that also shares an edge with all nodes within the clique so far is selected. 3. The process repeats until no nodes that share an edge with all nodes in the clique exist. 4. The clique is completed, all non-clique edges in the clique subgraph are removed, and a new unprocessed node with the greatest number of neighbors is selected and added to the current clique. 5. The process again repeats till the clique can not be expanded anymore. 6. The new clique is completed. Any singleton nodes remaining in the graph are marked as completed. The result can be separated into group cliques (in red) and singleton cliques (in green), which form the basis of the clique-based metrics used.

#### **Greedy Minimum Clique Cover Algorithm**

ALGORITHM Greedy Minimum Clique Cover

```
INPUT Graph G with vertices V and edges E: G(V, E)

GREEDY MINIMUM CLIQUE COVER( G(V, E) ):
	S ← {}
	while V is not empty
	for v ∈ V
		if degree(v) = Δ(G) then
```

v_0_ ← v

break

H ← {v_0_}

P ← ⋂_v ∈ H_ N(v)

```
		while P is not empty
			for p ∈ P
				if degree(p) = Δ(P) then
					H ← H ⋃ {p}
					break
```

P ← ⋂_v ∈ H_ N(v)

```
		V ← V \ H
		S ← S ⋃ {H}
	return S
```



Dau, H., Milenkovic, O., & Puleo, G. J. (2020). On the triangle clique cover and kt clique cover problems. *Discrete Mathematics*, *343*(1), 111627.

Karp, R. M. (2009). Reducibility among combinatorial problems. In *50 Years of Integer Programming 1958-2008: from the Early Years to the State-of-the-Art* (pp. 219-241). Berlin, Heidelberg: Springer Berlin Heidelberg.

**Supplementary Text 3. National Institute of Standards and Technology (NIST) Reference Material 1950 (SRM 1950) – Metabolites in frozen human plasma.**

The SRM 1950 unit contains vials containing approximately 1 mL of pooled human plasma, thawed on ice and vortex-mixed before use.

*Spike‑in standards*

A mixture of 126 metabolites was assembled from commercially available kits. The pool included six biochemical classes (amino acids, bile acids, organic acids, carnitines/acylcarnitines, catecholamines, and other metabolites). Stock solutions of individual compounds were supplied at a concentration of 1 mg/mL in water or methanol. Four mixed working solutions (low, medium, high, and super-high) were prepared by combining appropriate volumes of each individual stock with LC-MS-grade water/methanol. The resulting mixture contained 126 compounds at low, medium, high, and super-high concentrations, where the actual final concentrations ranged from 0.07 µM to 20 µM in the low mixture, 0.14–113 µM in the medium mixture, 0.28–80 µM in the high mixture, and 1.1–320 µM in the super‑high mixture.

*Preparation of spiked plasma samples*

All sample handling was performed on ice using low‑bind polypropylene tubes. Aliquots of SRM 1950 NIST human plasma (200 µL) were pipetted into pre-labeled tubes. Each of the four spike-in concentrations were added to separate plasma aliquots in triplicate. To achieve the target concentrations, 0.1 µL, 0.4 µL, 0.8 µL, or 1.6 µL of the mixed standard solution was added to each 200‑µL plasma aliquot (corresponding to the low, medium, high, and super‑high levels, respectively). Tubes were vortexed for 30 seconds, then incubated on ice for 10 minutes to allow equilibration. To monitor matrix effects, blank samples (prepared with water/methanol instead of plasma) and process blanks (methanol added to plasma without standards) were prepared in parallel.

*Plasma metabolite extraction*

Proteins in plasma samples (200 µL) were precipitated using 4x volume (800 µL) of chilled methanol (−20 °C). After vortex mixing for 1 min, samples were incubated on ice for 30 min and centrifuged at 15,000×g for 10 min at 4 °C. The clear supernatant (≈ 900 µL) was transferred to fresh tubes and dried under vacuum using a centrifugal evaporator. The residue was reconstituted in 200 µL of 5% acetonitrile containing 0.1% formic acid. Immediately before injection, samples were filtered through 0.22‑µm PTFE filters to remove particulates.

**Supplementary Text 4. Preparation of plant natural products.**

Purchased chemical standards **(Supplementary Table 2)** were dissolved in DMSO at 10 or 20 mM concentrations. A working pool was prepared by diluting each stock into reconstitution solvent to a final concentration of 150 ng/µL for each compound. Three serial dilutions of the pool 10, 20, and 40 ng/µL) were prepared, and the dilutions corresponded to 50, 100, and 200 ng of injected material when 5 or 10 µL were injected.

**Supplementary Text 5. Untargeted LC-MS/MS acquisition methods.**

Samples were analyzed on a Bruker timsTOF Pro instrument (Bruker, Bremen, Germany) equipped with an ultra‑high‑performance liquid chromatography (UHPLC) system (Waters ACQUITY), and separations were performed on an ACQUITY Premier HSS T3 1.8 µm, 50 x 2.1 mm C18 column (Part no. 186009467, Serial no. 03043322918640), where samples were acquired in a positive ESI mode. For the C18 method, 30 µL of extracted plasma was injected. Mobile phase A consisted of water with 0.1 % formic acid, and mobile phase B was methanol with 0.1 % formic acid. A linear gradient from 2 % to 98 % B was applied over 10 min at a flow rate of 0.4 mL min⁻¹, followed by a 2‑min wash at 98 % B and re‑equilibration. Column temperature was maintained at 40 °C.

timsTOF was operated in 4-dimensional data acquisition (PASEF) mode. Electrospray voltage was set to +4.5 kV (positive mode). Drying gas (nitrogen) was maintained at 8 L min⁻¹ and 220 °C. Ion mobility separation used a 1/K₀ scan range of 0.6–1.9 V s cm⁻² with a ramp time of 100 ms. Data was acquired over an *m/z* range of 50–1200 with an acquisition rate of 10 Hz. Internal calibrants were infused at the beginning and end of each batch to correct for mass and mobility drifts. Quality‑control (QC) samples were prepared by pooling equal aliquots of all spiked samples and injecting the QC every ten injections. Samples were injected in randomized order to minimize run-order effects. QC and blank injections ensured analytical stability and addressed carry-over issues.

#### **Supplementary Text 6. Parameters used to run MetaboScape.**

No minimum number of features for extraction, Intensity threshold: 1000 counts, minimum peak height: 6; Feature signal: Height; Recursive feature extraction was enabled with 75 datapoints; Mass range: 20-1500. No mass and IM calibrations were performed, as the publicly available datasets were incomplete and random due to a lack of calibration data.

#### **Supplementary Text 7. Parameters used to run MS-DIAL.**

Centroid input data, positive mode, standard *m/z* and RT ranges, MS1 mass tolerance at 0.01 Da, MS2 mass tolerance at 0.1 Da, minimum peak height (MS1) at 1000, minimum peak width at ≥5 scans, mass slice width at 0.1 Da, smoothing method being Linear‑Weighted Moving Average, smoothing level at 3; alignment MS1 tolerance at 0.01 Da, alignment RT tolerance at 0.1  min without any gap filling & missing‑value handling, and no feature cleanup or blank filters were applied.

#### **Supplementary Text 8. Parameter file used to run MSDIAL.**

# Project information

```
MS-DIAL version number: 5.5.250221
Project folder path:
Project file path:
MS1 data type: Centroid
MS2 data type: Centroid
Ion mode: Positive
Target omics: Metabolomics
Ionization: ESI
Machine category: LCIMMS
Instrument type:
Instrument:
Authors:
License:
Comment:
Msp file path:
Lbm file path:
Text DB file path:
Isotope text DB file path:
Compounds library file path for target detection:
Compounds library file path for RT correction:
Searched adduct ions:

# FileID ClassName

# FileID AnalysisFileType

# Classname order

# Classname ColorBytes

# Export
Export spectra file format: msp
Export spectra type: deconvoluted
Mat file export folder path:
Export folder path:
Height matrix export: True
Normalized height matrix export: False
Representative spectra export: False
Peak ID matrix export: False
Retention time matrix export: False
Mass matrix export: False
MSMS included matrix export: False
Unique mass matrix export: False
Peak area matrix export: False
Parameter export: False
GNPS export: False
Molecular networking export: False
SN matrix export: False
Export as mztabM format: False

# Process parameters
Process option: All
Number of threads: 16

# Feature detection parameters
Smoothing method: LinearWeightedMovingAverage
Smoothing level: 3
Minimum peak height: 1000
Minimum peak width: 5
Average peak width: 30
Mass slice width: 0.1
Retention time begin: 0
Retention time end: 100
MS1 mass range begin: 0
MS1 mass range end: 2000
MS2 mass range begin: 0
MS2 mass range end: 2000
MS1 tolerance for centroid: 0.01
MS2 tolerance for centroid: 0.1
Accuracy type: IsAccurate
Max charge number: 2
Considering Br and Cl for isotopes: False
Exclude mass list:
Max isotopes detected in ms1 spectrum: 2

# Deconvolution
Sigma window value: 0.5
Amplitude cut off: 0
Keep isotope range: 5
Exclude after precursor: True
Keep original precursor isotopes: False
Is do andromeda ms2 deconvolution: False
Andromeda delta: 100
Andromeda max peaks: 12
Target CE: 0

# Annotation parameter

# Retention index dictionary information


# Alignment parameters
Alignment reference file ID: 0
Retention time tolerance for alignment: 0.1
Retention time factor for alignment: 0.5
Spectrum similarity tolerance for alignment: 0.8
Spectrum similarity factor for alignment: 0.5
MS1 tolerance for alignment: 0.01
MS1 factor for alignment: 0.5
Force insert peaks in gap filling: False

# Filtering
Peak count filter: 0
N percent detected in one group: 0
Remove feature based on peak height fold-change: False
Blank filtering: SampleAveOverBlankAve
Sample max / blank average: 5
Sample average / blank average: 5
Keep reference matched metabolites: False
Keep suggested metabolites: False
Keep removable features and assigned tag for checking: False
Replace true zero values with 1/2 of minimum peak height over all samples: False

# Retention time correction
Execute RT correction: False
RT correction with smoothing for RT diff: False
User setting intercept: 0
RT diff calc method: SampleMinusSampleAverage
Interpolation method: Linear
Extrapolation method (begin): UserSetting
Extrapolation method (end): LastPoint
Internal standards for RT alignment:

# Isotope tracking setting
Tracking isotope label: False
Set fully labeled reference file: False
Non labeled reference ID: 0
Fully labeled reference ID: 0
Isotope tracking dictionary ID: 0

# CorrDec settings
CorrDec execute: True
CorrDec MS2 tolerance: 0.01
CorrDec minimum MS2 peak height: 1000
CorrDec minimum number of detected samples: 3
CorrDec exclude highly correlated spots: 0.9
CorrDec minimum correlation coefficient (MS2): 0.7
CorrDec margin 1 (target precursor): 0.2
CorrDec margin 2 (coeluted precursor): 0.1
CorrDec minimum detected rate: 0.5
CorrDec minimum MS2 relative intensity: 2
CorrDec remove peaks larger than precursor: True

# IMMS specific parameters
Drift time begin: 0
Drift time end: 2000
Accumulated RT ragne: 0.2
Accumulate MS2 spectra: False
Drift time alignment tolerance: 0.02
Drift time alignment factor: 0.5
Ion mobility type: Tims
All calibrant data imported: False

# File ID CCS coefficients
DataBaseID:
AnnotationMethod:
MassRangeBegin: 0.00000
MassRangeEnd: 2000.00000
RtTolerance: 100.00
RiTolerance: 100.000
CcsTolerance: 50.00
Ms1Tolerance: 0.05000
Ms2Tolerance: 0.10000
RelativeAmpCutoff: 0.000
AbsoluteAmpCutoff: 0
WeightedDotProductCutOff: 0.100
SimpleDotProductCutOff: 0.100
ReverseDotProductCutOff: 0.100
MatchedPeaksPercentageCutOff: 0.000
AndromedaScoreCutOff: 0.100
TotalScoreCutoff: 0.800
MinimumSpectrumMatch: 0.000
IsUseTimeForAnnotationFiltering: False
IsUseTimeForAnnotationScoring: False
IsUseCcsForAnnotationFiltering: False
IsUseCcsForAnnotationScoring: False

DataBaseID: MS-FINDER
```

#### **Supplementary Text 9. Parameters used to run MZmine.**

MS1 noise level 1.0E2, MS2 noise level 2.0E1; Chromatogram builder (ADAP): *m/z* tolerance 0.0050 *m/z* or 20.0 ppm, Minimum consecutive scans 4; Minimum intensity for consecutive scans 1.0E3, Minimum absolute height 1.0E3; Local minimum feature resolver (RT dimension): Chromatographic threshold 90.0%, Minimum absolute height 1.0E3, Minimum ratio of peak top/edge 1.80, Peak duration range 0.00 - 1.51, Minimum search range RT (absolute): 0.050 ; Ims expander: *m/z* tolerance 0.0050 *m/z* or 20.0 ppm; Local minimum feature resolver (Mobility dimension): Chromatographic threshold 80.0%, Minimum absolute height 1.0E3, Minimum ratio of peak top/edge 1.80, Peak duration range 0.00 - 20.00, Minimum search range Mobility (absolute) 0.010; 13C isotope filter: *m/z* tolerance (intra-sample) 0.0015 *m/z* or 3.0 ppm, Retention time tolerance 0.04 minutes, Mobility tolerance 0.008, Maximum charge 2, Representative isotope Most intense; Ion identity networking: *m/z* tolerance (intra-sample) 0.0015 *m/z* or 3.0 ppm; Join aligner: *m/z* tolerance (sample-to-sample) 0.0040 *m/z* or 8.0 ppm, Weight for *m/z* 3, Retention time tolerance 0.10 minutes, Weight for RT 1, Mobility tolerance 0.01, Mobility weight 1.000; Feature list rows filter: At least 1 aligned sample, Never remove rows with MS2; Feature finder (multithreaded): Intensity tolerance 20.0%, *m/z* tolerance (sample-to-sample) 0.0050 *m/z* or 20.0 ppm, Retention time tolerance 0.10 minutes, Minimum scans 2; Duplicate peak filter: *m/z* tolerance 0.0008 *m/z* or 1.5 ppm, RT tolerance 0.04 minutes, Mobility tolerance 0.008; Correlation grouping (metaCorrelate): RT tolerance 0.06 minutes, Minimum feature height 0.0E0, Intensity threshold for correlation 5.0E2. QC/blank filtering or feature presence threshold was applied. A detailed batch file (.mzbatch) is provided as a supplementary file **(Supplementary File 2)**.

### **Supplementary Tables**

| **Dataset Type** | **Dataset ID** | **File Counts** | **Type of dataset** | **LC methods** | **Software used by Authors** | **Paper** |
| --- | --- | --- | --- | --- | --- | --- |
| *Ground-Truth (Public)* | [MSV000098263](https://massive.ucsd.edu/ProteoSAFe/dataset.jsp?task=5094dadc7b754e98b01c718ca09dac84) | 748 | Drugs: Crude standards of reframe 4723 drug mixtures were analyzed on a TimsTOF Pro2 | C18 Reversed phase, U-Flow (UHPLC) | MetaboScape2025 | [Link](https://pmc.ncbi.nlm.nih.gov/articles/PMC11482764/#S2) |
| *Ground-Truth (Internal)* | Internal ID | 80 | NIST® SRM® 1950 human blood plasma was spiked with a mixture of 120 metabolites, at four different concentrations, all run together with native plasma as well. | C18 Reversed phase, U-Flow (UHPLC) | NA | NA |
| *Ground-Truth (Internal)* | Internal ID | 57 | 45 Plant Natural Products ran individually, at three different on-column amounts (50, 100, and 200 ng) | C18 Reversed phase, U-Flow (UHPLC) | NA | NA |
| Public | [MSV000090327](https://massive.ucsd.edu/ProteoSAFe/dataset.jsp?accession=MSV000090327) | 46 | Plant: Ethanolic extracts of piper plants (leaves, fruits) | C18 Reversed phase, U-Flow (UHPLC) | MZmine 3.3.0 | [Link](https://pmc.ncbi.nlm.nih.gov/articles/PMC10496610/) |
| Public | [MSV000091642](https://massive.ucsd.edu/ProteoSAFe/dataset.jsp?task=938ac19eec3e4c5bbcb64ab765eb6e1b) | 3 | Animal: Sheep brain samples extracted with methyl-tert-butyl ether (Matyash protocol) | HILIC | MZmine 3.3.0 | [Link](https://pmc.ncbi.nlm.nih.gov/articles/PMC10496610/) |
| Public | [MSV000095813](https://massive.ucsd.edu/ProteoSAFe/dataset.jsp?task=d3a4caa6ec9f4d4f81ba235984d054c7) | 16 | Microbial (*Trichoderma reese*i ) | C18 Reversed phase, U-Flow (UHPLC) | MZmine 4.0.3 | [Link](https://pubmed.ncbi.nlm.nih.gov/40069746/) |
| Public | [MSV000097967](https://massive.ucsd.edu/ProteoSAFe/dataset.jsp?task=cc2496604f0149e2b685f7c721712ffb) | 62 | Gut Microbiota: Enteromix model of the human large intestine | C18 Reverse phase, U-Flow (UHPLC) | MZmine 4.4.3  MetaboScape | [Link](https://www.biorxiv.org/content/10.1101/2025.09.25.678496v1) |
| Public | [MSV000097015](https://massive.ucsd.edu/ProteoSAFe/dataset.jsp?task=7ea04cbbd9b04840a3d3dcb9c5c1b846) | 3 | Human: Serum for glycosphingolipidome profiling | C18 Reversed phase, U-Flow (UHPLC) | MetaboScape 2021b | [Link](https://pmc.ncbi.nlm.nih.gov/articles/PMC12084332/) |
| Public | [MSV000096291](https://massive.ucsd.edu/ProteoSAFe/dataset.jsp?task=a9da614e256546b3a63c469dd7965dac) | 161 | Standards: Bile acid fragmentation dataset | C18 Reversed phase, U-Flow (UHPLC) | MZmine 4.2.0 | [Link](https://www.biorxiv.org/content/10.1101/2025.03.04.641505v1.full) |
| Public | [MSV000096189](https://massive.ucsd.edu/ProteoSAFe/dataset.jsp?task=a671eaf06a4440ea9d555c7a43ba8131) | 21 | Mice: Macrophages, WT murine bone marrow | HILIC, UPLC | otofContro, timsControl, Skyline | [Link](https://www.sciencedirect.com/science/article/pii/S000326702500399X) |
| Public | [ST002402](https://www.metabolomicsworkbench.org/data/DRCCMetadata.php?Mode=Study&StudyType=&StudyID=ST002402) | 34 | Human: NIST 1950 SRM plasma, 1951 SRM serum, blood, venous, and finger-prick dried blood spots- chose the smallest 1-month treated dataset | C18, UHPLC | MetaboScape 2021a | [Link](https://www.nature.com/articles/s41467-023-36520-1) |
| Public | [MSV000084402](https://massive.ucsd.edu/ProteoSAFe/dataset.jsp?task=36fea50f5e7b4a049d336f28c5884ff9) | 1 | Human: NIST 1950 SRM plasma single sample dataset | C18, UHPLC | MetaboScape | [Link](https://www.nature.com/articles/s41592-020-0933-6) |
| Public | [MTBLS12332](https://www.ebi.ac.uk/metabolights/editor/MTBLS12332) | 313 | Human: HepG-2 cells treated with 11 test compounds | C18 Reversed phase, U-Flow (UPLC) | MetaboScape | [Link](https://link.springer.com/article/10.1007/s11306-025-02309-0#Sec12) |

#### **Supplementary Table 1. List of all datasets used for the study.**

| **Compound Name** | **Category** | **SMILES** | **Molecular Formula** | **Molecular Weight (Da)** | **XLogP** |
| --- | --- | --- | --- | --- | --- |
| 10-gingerol | Phenylpropanoids | CCCCCC=CC(=O)CCCC1=CC=CC=C1O | C21H32O4 | 332.48 | 5.3 |
| 4-hydroxybenzyl alcohol | Phenolics | OCc1ccc(O)cc1 | C7H8O2 | 124.14 | 0.2 |
| aloperine | Alkaloids | C1CN2CCC(C3=CC=CC=C3C2C1)N | C15H24N2 | 232.36 | 1.6 |
| caftaric acid | Phenylpropanoids | O=C(O)C=CC1=CC(=C(C=C1)O)OCC(=O)O | C13H12O9 | 312.23 | 0.1 |
| cannabidivarin | Phenylpropanoids | CCC1=C(C=C2C(=C1)C(CCC(=O)C)O2)C | C19H26O2 | 286.41 | 5.4 |
| catharanthine | Alkaloids | COC1=CC2=C(C=C1)C=CN(C3=C2C=CC(=O)N3C)C4CC4 | C21H24N2O2 | 336.43 | 2.8 |
| chelidonine | Alkaloids | O=C1COC2=C(C1)C=CC=C2N3CC=CC4=C3C=CC(=O)O4 | C20H19NO5 | 353.37 | 1.5 |
| cinchonidine | Alkaloids | C=1C=CC=CC=1[C@H]2O[C@H]3N=C(C=C2N3)C(C)=C | C19H22N2O | 294.39 | 2.7 |
| colchicine | Alkaloids | COC1=CC=CC2=C1C(=C(N(C2=O)C)C3=CC=CC=C3)OC | C22H25NO6 | 399.44 | -0.1 |
| coumarin | Phenylpropanoids | O=C1C=CC=CO1C2=CC=CC=C2 | C9H6O2 | 146.14 | 1.4 |
| cryptotanshinone | Diterpenoid Quinone | C[C@H]1COC2=C1C(=O)C(=O)C3=C2C=CC4=C3CCCC4(C)C | C19H20O3 | 296.3 | 3 |
| dehydroevodiamine | Alkaloids | COC1=CC2=C(C=C1)C=CN(C3=CC=CC=C3)C2=C | C19H17N3 | 287.36 | 2.1 |
| dehydronuciferine | Alkaloids | COC1=CC2=C(C=C1)C=CN(C)C3=CC4=C(C=C3C2)OCO4 | C19H15NO2 | 289.33 | 4.5 |
| dihydrochelerythrine | Alkaloids | COC1=CC2=C(C=C1)C=C[N+]3=C2C=CC(=C3)C4=CC(=C(C=C4)OC)OC | C21H19NO4+ | 349.38 | 2.1 |
| eriodictyol | Flavonoids | O=C1C(OC)=C(O)C(=O)C2=CC(O)=CC(O)=C2C1 | C15H12O6 | 288.25 | 2 |
| glabridin | Flavonoids | O=C1C(O)=C(O)C(=O)C2=CC(=C(C=C2C1C3=CC=CC=C3OC)OC)OC | C20H20O4 | 324.37 | 3.9 |
| hederagenin | Triterpenoid | C[C@]12CC[C@@H]([C@@]([C@@H]1CC[C@@]3([C@@H]2CC=C4[C@]3(CC[C@@]5([C@H]4CC(CC5)(C)C)C(=O)O)C)C)(C)CO)O | C30H48O4 | 472.7 | 4.8 |
| hordenine | Alkaloids | Oc1ccc(CCN(C)C)cc1 | C10H15NO | 165.23 | 2.1 |
| hydroxytyrosol acetate | Phenolics | CC(=O)OC1=CC(O)=CC(O)=C1 | C9H10O4 | 182.17 | -0.1 |
| hydroxy-α-sanshool | Phenylpropanoids | CCC=CC=CC(C=O)NC(=O)C=CC=CC=CC | C14H23NO2 | 237.34 | 3.4 |
| indole-3-carbinol | Alkaloids | OCC1=CNC2=CC=CC=C12 | C9H9NO | 147.17 | 1.1 |
| karanjin | Flavonoids | O=C1C=C2C(=O)OC3=CC=CC=C3C2=CN1C4=CC=CC=C4 | C18H12O4 | 292.29 | 3.8 |
| lotusine | Alkaloids | COc1ccc2c(c1OC)C=C[N+](C)C3=C2C=CC(=O)O3 | C19H18NO4+ | 320.35 | 3.1 |
| lupinine | Alkaloids | C1CN2CCCC2C(C1)O | C10H19NO | 169.26 | 1.2 |
| madecassic acid | Triterpenoid | C[C@@H]1CC[C@@]2(CC[C@@]3(C(=CC[C@H]4[C@]3(C[C@H]([C@@H]5[C@@]4(C[C@H]([C@@H]([C@@]5(C)CO)O)O)C)O)C)[C@@H]2[C@H]1C)C)C(=O)O | C30H48O6 | 504.7 | 4.7 |
| maltol | Flavonoids | O=C1C=CC(O)=C(O1)C | C6H6O3 | 126.11 | 0.4 |
| methyl gallate | Phenolics | COC(=O)C1=CC(=C(C(=C1)O)O)O | C8H8O5 | 184.15 | 0.9 |
| neobavaisoflavone | Flavonoids | OC1=C(C=CC(=C1)C2=CC(=O)C3=C(C2=O)C=CC=C3)C | C20H12O4 | 316.31 | 4.4 |
| oxindole | Alkaloids | O=C2NC1=CC=CC=C1C2 | C8H7NO | 133.2 | 1.2 |
| oxysophocarpine | Alkaloids | O=C1CN2CCC=C(C2C(C1)O)C3=CC=CC=C3 | C15H22N2O2 | 262.35 | 1.2 |
| piperine | Alkaloids | OC(=O)C=CC1=CC=C(N2CCCCC2=O)C=C1 | C17H19NO3 | 285.34 | 3.5 |
| piperlongumine | Alkaloids | O=C1C=CC2=CC=CC=C2C1=CC(=O)N3CCCCC3 | C17H19NO5 | 317.34 | 2.1 |
| plumbagin | Phenolics | CC1=CC(=O)C2=C(C1=O)C=CC=C2O | C11H8O3 | 188.2 | 2 |
| protocatechualdehyde | Phenolics | O=CC1=CC(=CC(=C1)O)O | C7H6O3 | 138.07 | 1.3 |
| psoralen | Coumarins | O=C1OC2=CC=CC=C2C3=C1C=CC=C3 | C11H6O3 | 186.17 | 2.3 |
| sanguinarine | Alkaloids | COC1=CC2=C(C=C1)C=CC3=C2C=CC(=[N+](C)C=C3)C | C20H14NO4+ | 314.33 | 4.4 |
| sinomenine | Alkaloids | COC1=CC2=C(C=C1O)C(CC3=C2N(C)C=C(C3)C)=O | C19H23NO4 | 329.39 | 2.2 |
| synephrine | Alkaloids | OC(C1=CC=CC=C1)C(N)C | C9H13NO2 | 167.21 | -0.6 |
| trans-zeatin | Purines | C=C(N)CNc1nc(N)nc2c1[nH]cn2 | C10H13N5O | 219.24 | 0.7 |
| tropine | Alkaloids | C1CN2CCCCC2C1O | C8H15NO | 141.21 | 0.8 |
| tropinone | Alkaloids | O=C1CC2CCCN2C1 | C8H13NO | 139.19 | 0.3 |
| ursolic acid | Triterpenoid | C[C@@H]1CC[C@@]2(CC[C@@]3(C(=CC[C@H]4[C@]3(CC[C@@H]5[C@@]4(CC[C@@H](C5(C)C)O)C)C)[C@@H]2[C@H]1C)C)C(=O)O | C30H48O3 | 456.7 | 6.8 |
| visnagin | Phenylpropanoids | O=C1OC2=CC=CC=C2C(=C1C)C3=CC=CC=C3 | C13H10O3 | 214.22 | 2.3 |
| withaferin A | Steroidal Lactone | CC1=C(C(=O)O[C@H](C1)[C@@H](C)[C@H]2CC[C@@H]3[C@@]2(CC[C@H]4[C@H]3C[C@@H]5[C@]6([C@@]4(C(=O)C=C[C@@H]6O)C)O5)C)CO | C28H38O6 | 470.6 | 2.7 |
| xanthotoxol | Coumarins | O=C1OC2=CC=CC=C2C3=C1C=CC(O)=C3 | C11H6O4 | 202.17 | 1.6 |

#### **Supplementary Table 2. List of plant-derived natural products spike-in ground-truth data with 45 compounds injected at three on column amounts (50, 100, and 200 ng).**

| **Adduct** | **Ion mode** | **Charge** | **Mass multiplier** | **Correction mass** | **Priority** | **Primary** |
| --- | --- | --- | --- | --- | --- | --- |
| [M+2H-H2O]2+ | positive | 2 | 0.5 | -15.99491462 | 0 | 1 |
| [M+2H-2H2O]2+ | positive | 2 | 0.5 | -34.0054793 | 0 | 1 |
| [M+2H]2+ | positive | 2 | 0.5 | 2.015650064 | 0.2 | 1 |
| [M+H+NH4]2+ | positive | 2 | 0.5 | 19.04219916 | 0 | 1 |
| [M+H+Na]2+ | positive | 2 | 0.5 | 23.99759431 | 0 | 1 |
| [M+H+K]2+ | positive | 2 | 0.5 | 39.97153151 | 0 | 1 |
| [M+2Na]2+ | positive | 2 | 0.5 | 45.97953856 | 0 | 1 |
| [M+Na+K]2+ | positive | 2 | 0.5 | 61.95347576 | 0 | 1 |
| [M+2K]2+ | positive | 2 | 0.5 | 77.92741296 | 0 | 1 |
| [M+H]+ | positive | 1 | 1 | 1.007825032 | 0.8 | 1 |
| [M+H-C6H10O5]+ | positive | 1 | 1 | -161.0449984 | 0 | 1 |
| [M+H-CH4O3]+ | positive | 1 | 1 | -63.00821896 | 0 | 0 |
| [M+H-C2H4O2]+ | positive | 1 | 1 | -59.01330434 | 0 | 0 |
| [M+H-C4H8]+ | positive | 1 | 1 | -55.05477523 | 0 | 1 |
| [M+H-C2H6O]+ | positive | 1 | 1 | -45.03403978 | 0 | 0 |
| [M+H-CH2O2]+ | positive | 1 | 1 | -44.99765427 | 0 | 0 |
| [M+H-C2H7N]+ | positive | 1 | 1 | -44.0500242 | 0 | 0 |
| [M+H-CO2]+ | positive | 1 | 1 | -42.98200421 | 0 | 0 |
| [M+H-C3H6]+ | positive | 1 | 1 | -41.03912516 | 0 | 1 |
| [M+H-C2H2O]+ | positive | 1 | 1 | -41.00273965 | 0 | 1 |
| [M+H-CH4O]+ | positive | 1 | 1 | -31.01838972 | 0 | 0 |
| [M+H-CH5N]+ | positive | 1 | 1 | -30.03437413 | 0 | 1 |
| [M+H-C2H4]+ | positive | 1 | 1 | -27.0234751 | 0 | 0 |
| [M+H-CO]+ | positive | 1 | 1 | -26.98708959 | 0 | 1 |
| [M+H-NH3]+ | positive | 1 | 1 | -16.01872407 | 0 | 1 |
| [M+H-CH3]+ | positive | 1 | 1 | -14.01565006 | 0 | 1 |
| [M+H-H2O]+ | positive | 1 | 1 | -17.00273965 | 0.5 | 1 |
| [M+H-2H2O]+ | positive | 1 | 1 | -35.01330434 | 0 | 1 |
| [M+H-3H2O]+ | positive | 1 | 1 | -53.02386902 | 0 | 1 |
| [M+NH4]+ | positive | 1 | 1 | 18.03437413 | 0.4 | 1 |
| [M+K-H2O]+ | positive | 1 | 1 | 20.9531418 | 0 | 1 |
| [M+Na]+ | positive | 1 | 1 | 22.98976928 | 0.7 | 1 |
| [M+H+CH3OH]+ | positive | 1 | 1 | 33.03403978 | 0 | 1 |
| [M+K]+ | positive | 1 | 1 | 38.96370649 | 0.2 | 1 |
| [2M+H-H2O]+ | positive | 1 | 2 | -17.00273965 | 0 | 1 |
| [2M+H]+ | positive | 1 | 2 | 1.007825032 | 0.4 | 1 |
| [2M+NH4]+ | positive | 1 | 2 | 18.03437413 | 0.5 | 1 |
| [2M+Na]+ | positive | 1 | 2 | 22.98976928 | 0.3 | 1 |
| [2M+H+CH3OH]+ | positive | 1 | 2 | 33.03403978 | 0 | 1 |
| [2M+K]+ | positive | 1 | 2 | 38.96370649 | 0 | 1 |
| [M-2H]2- | negative | -2 | 0.5 | -2.015650064 | 0.5 | 1 |
| [M-H-H2O]- | negative | -1 | 1 | -19.01838972 | 0.5 | 1 |
| [M-H]- | negative | -1 | 1 | -1.007825032 | 0.8 | 1 |
| [M-H-C6H10O5]- | negative | -1 | 1 | -163.0606485 | 0 | 1 |
| [M-H-CO2]- | negative | -1 | 1 | -44.99765427 | 0 | 1 |
| [M-H-CH3]- | negative | -1 | 1 | -16.03130013 | 0 | 1 |
| [M-2H+Na]- | negative | -1 | 1 | 20.97411922 | 0 | 1 |
| [M+Cl]- | negative | -1 | 1 | 34.96885268 | 0 | 1 |
| [M-2H+K]- | negative | -1 | 1 | 36.94805642 | 0 | 1 |
| [M-H+FA]- | negative | -1 | 1 | 44.99765427 | 0 | 1 |
| [2M-H]- | negative | -1 | 2 | -1.007825032 | 0 | 1 |
| [2M-H+FA]- | negative | -1 | 2 | 44.99765427 | 0 | 1 |
| [3M-H]- | negative | -1 | 3 | -1.007825032 | 0 | 1 |

#### **Supplementary Table 3. List of adducts used for feature calling in all three software tools (MZmine, MS-DIAL, and MetaboScape).**
